# Design, Construction, and Functional Characterization of a tRNA Neochromosome in Yeast

**DOI:** 10.1101/2022.10.03.510608

**Authors:** Daniel Schindler, Roy S.K. Walker, Shuangying Jiang, Aaron N. Brooks, Yun Wang, Carolin A. Müller, Charlotte Cockram, Yisha Luo, Alicia García, Daniel Schraivogel, Julien Mozziconacci, Benjamin A. Blount, Jitong Cai, Lois Ogunlana, Wei Liu, Katarina Jönsson, Dariusz Abramczyk, Eva Garcia-Ruiz, Tomasz W. Turowski, Reem Swidah, Tom Ellis, Francisco Antequera, Yue Shen, Conrad A. Nieduszynski, Romain Koszul, Junbiao Dai, Lars M. Steinmetz, Jef D. Boeke, Yizhi Cai

## Abstract

Here we report the design, construction and characterization of a tRNA neochromosome, a designer chromosome that functions as an additional, *de novo* counterpart to the native complement of *Saccharomyces cerevisiae*. Intending to address one of the central design principles of the Sc2.0 project, the ∼190 kb tRNA neochromosome houses all 275 relocated nuclear tRNA genes. To maximize stability, the design incorporated orthogonal genetic elements from non-*S. cerevisiae* yeast species. Furthermore, the presence of 283 *rox* recombination sites enable an orthogonal SCRaMbLE system capable of adjusting tRNA abundance. Following construction, we obtained evidence of a potent selective force once the neochromosome was introduced into yeast cells, manifesting as a spontaneous doubling in cell ploidy. Furthermore, tRNA sequencing, transcriptomics, proteomics, nucleosome mapping, replication profiling, FISH and Hi-C were undertaken to investigate questions of tRNA neochromosome behavior and function. Its construction demonstrates the remarkable tractability of the yeast model and opens up new opportunities to directly test hypotheses surrounding these essential non-coding RNAs.

**Highlights:**

- *De novo* design, construction and functional characterization of a neochromosome containing all 275 nuclear tRNA genes of *Saccharomyces cerevisiae*.
- Increasing the copy number of the 275 highly expressed tRNA genes causes cellular burden, which the host cell likely buffers either by selecting for partial tRNA neochromosome deletions or by increasing its ploidy.
- The tRNA neochromosome can be chemically extracted and transformed into new strain backgrounds, enabling its transplantation into multi-synthetic chromosome strains to finalize the Sc2.0 strain.
- Comprehensive functional characterization does not pinpoint a singular cause for the cellular burden caused by the tRNA neochromosome, but does reveal novel insights into its tRNA and structural chromosome biology.

## Introduction

Synthetic genomics is a nascent field of research that leverages advances in large-scale *de novo* DNA synthesis with rational design-based approaches of synthetic biology. Progressive advances in this field have led to the construction of viral genomes (Cello *et al*., 2002; Smith *et al*., 2003), bacterial genomes (Fredens *et al*., 2019; Gibson *et al*., 2010) and eukaryotic chromosomes (Annaluru *et al*., 2014; Richardson *et al*., 2017; Schindler *et al*., 2018). Developments in eukaryotes include work of the international synthetic yeast genome project (Sc2.0), an international endeavor which aims to re-design and chemically synthesize a designer version of the *Saccharomyces cerevisiae* genome. Unique design changes, such as the introduction of loxPsym sites for SCRaMbLE, the ‘freeing-up’ of the TAG stop codon for future reassignment and the removal of pre-mRNA and pre-tRNA introns (Dymond *et al*., 2011; Richardson *et al*., 2017; Shen *et al*., 2016), enable research into fundamental questions of biological significance which would otherwise not be possible through conventional means.

Sc2.0 is framed around three central design principles (Dymond *et al*., 2011; Richardson *et al*., 2017) which emphasize near wild-type fitness, the introduction of genetic flexibility to facilitate future research and the removal of potentially unstable elements such as tRNA genes (tDNAs) and transposons. While there are 61 codons specifying the 20 canonical amino acids, there are only 42 isoacceptor types of tRNA in *S. cerevisiae*, reflecting the fact that many tRNAs can recognize more than one codon through wobble interactions. As expected, a strong correlation exists between tRNA gene copy number and usage in translation (Percudani *et al*., 1997). Since tRNA genes are collectively but not necessarily individually essential to cell viability, Sc2.0 design specifications dictate the relocation of all tDNAs onto a dedicated synthetic chromosome (termed a tRNA neochromosome), so that the overall abundance of tRNA isoacceptor gene copies in the final synthetic strain matches that of the wild-type genome (Richardson *et al*., 2017). Furthermore, design specifications include the removal of tRNA introns from those copies that contain them. In this study, we report the design, construction and characterization of a tRNA neochromosome, a fully synthetic, designer chromosome which exists as an additional copy to the sixteen native chromosomes of *S. cerevisiae*.

tRNA molecules are fundamental adaptor molecules involved in translation, serving as a central link between the three-base codons of mRNA and the amino acids which form proteins. The genome of *S. cerevisiae* houses a total of 275 nuclear tRNA genes, including one pseudogene, dispersed throughout the genome (Hani and Feldmann, 1998). tRNA genes are transcribed by RNA polymerase III (RNAPIII) and associated transcription factors by conserved dual promoter elements (A and B sequence blocks) located internally to the tDNA (Geiduschek and Kassavetis, 2001). Due to their central role in biology, tRNA molecules are evolutionarily conserved and essential to all known free-living organisms.

tDNAs are known genomic instability hotspots for at least two reasons. Polar replication fork collisions between components associated with heavily transcribed tRNA genes cause replication fork stall events, leading to DNA breakage and chromosomal instability (Admire *et al*., 2006; Deshpande and Newlon, 1996; Hamperl *et al*., 2017). Furthermore, highly repetitive retrotransposons of up to 6 kb in length as well their solo LTR derivatives of 300-400 bp preferentially associate with regions upstream of tDNAs (Ji *et al*., 1993; Mularoni *et al*., 2012), and thus can act as hotspots for chromosomal recombination (Dunham *et al*., 2002; Mieczkowski *et al*., 2006). Our rationale is that removing repetitive elements and relocating tRNA genes onto a dedicated synthetic chromosome will lead to a synthetic yeast genome that is less susceptible to structural rearrangement and isolate any instability caused by their presence (Dymond *et al*., 2011). The tRNA neochromosome also serves as a unique chassis to systematically explore tRNA genetics through the Dre-*rox* SCRaMbLE system, providing a technological platform to facilitate research into tRNA and chromosomal biology, and demonstrate design-driven engineering biology approaches to enable the construction of future neochromosomes.

## Results

### Design of the tRNA Neochromosome

In contrast to other synthetic chromosomes of the Sc2.0 project, the tRNA neochromosome has no native counterpart in the yeast genome. As no pre-existing chromosome is available to serve as a template, the tRNA neochromosome was designed and built *de novo*. To rationalize the design process, we arranged each element of the tRNA neochromosome into defined layers of structural hierarchy (Fig. 1 A). The smallest of these layers consisted of 275 tRNA ‘cassettes’ (∼680 bp) concatenated into 16 tRNA ‘arrays’ (2.6 kb to 23.3 kb; Tab. S3 and S5), each designed to house tRNA genes from an *S. cerevisiae* native chromosome (Fig. 1A and Tab. S3). These tRNA arrays in turn comprised ‘mega-arrays’ consisting of one to three tRNA arrays orientated in tandem so as to be transcribed and replicated in the same direction (Fig. 1B). Following linearization and release of functional telomere seed sequences through deployment of the telomerator system (Mitchell and Boeke, 2014), the circular tRNA neochromosome was converted into a true linear chromosome.

#### Box 1 Summary of tRNA neochromosome design features.

##### Structural Hierarchy

The tRNA neochromosome consists of four levels of hierarchy. These are (from smallest to largest): tRNA cassettes, tRNA arrays, mega-arrays and the overall structure of the tRNA neochromosome (Fig. 1A).

##### Maximizing Stability

###### Orthogonality

To reduce homology to the host genome, the tRNA neochromosome consisted of genetic elements recovered from yeast species that are distinct from, but closely related to, *S. cerevisiae* where possible.

###### Reducing Replication Stress

The replication profile was designed to minimize replication stress associated with polar replication fork collisions. tRNA genes point away from each replication fork. DNA replication was designed to terminate at defined regions.

##### Automation of tDNA Flanking Sequence Assignment

Although Sc2.0 developed the BioStudio software to design synthetic yeast chromosomes (Richardson *et al*., 2017), the unique structure of the tRNA neochromosome necessitated the development of custom software. An algorithm and scripts (Fig. S1) were used to assign tDNAs to flanking sequences recovered from *A. gossypii* and *E. cymbalariae*.

##### Dre-*rox* Recombination System

*rox* recombination sites were incorporated to flank each tRNA cassette and functional genetic elements to provide a future system to study tRNA genetics through copy number adjustment.

##### Removal of tRNA Introns

The original Sc2.0 design objectives included the intent to explore the essentiality of tRNA introns, and thus all tRNA introns were removed.

**Fig. 1.**
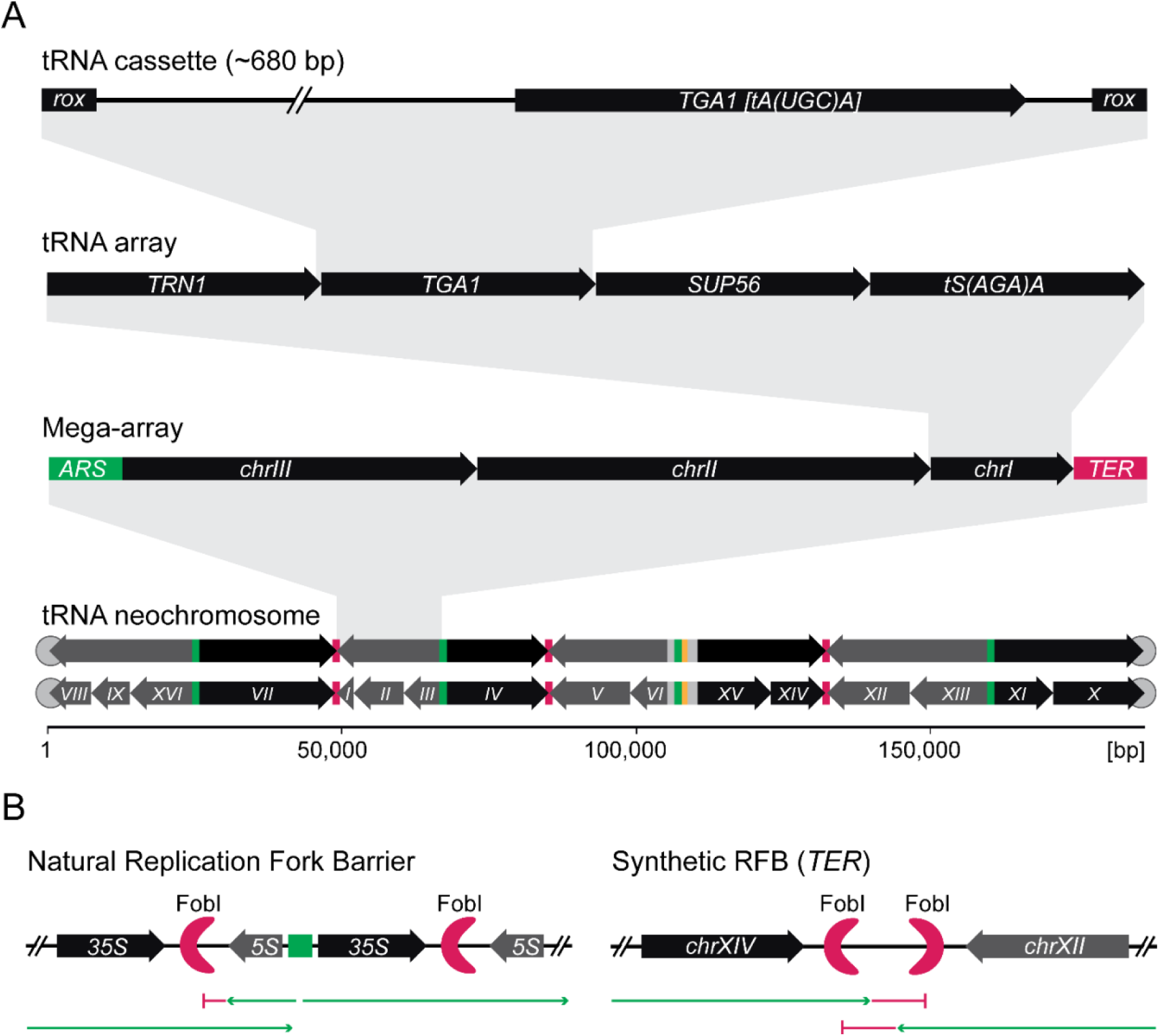
Design and hierarchy of the tRNA neochromosome. **(A)** The smallest level of tRNA neochromosome hierarchy includes the individual tRNA cassettes consisting of orthogonal 500 bp 5’ and 40 bp 3’ sequences flanking each *S. cerevisiae* tRNA coding region (intronless for tRNA genes containing introns). These are in turn flanked by *rox* recombination sites. These tRNA cassettes are next are next incorporated into a chromosome-specific tRNA array in which transcription of all tRNAs point in the same direction. tRNA arrays are in turn ordered into mega-arrays of approximately even size, with DNA replication and transcription orientated to proceed in the same direction. Autonomous replicating sequences (*ARS*) are indicated in green, synthetic bidirectional termination sites (*TER*) are indicated in red and pRS413 centromere region is indicated in orange. The tRNA neochromosome itself consists of 275 tRNA cassettes ordered in 16 tRNA arrays which then comprise 8 mega-arrays (detailed data in supporting information table S5). The final tRNA neochromosome can be subsequently equipped with telomeres indicated by gray circles (Mitchell and Boeke, 2014) to engineer sites of linearization. Roman numbers indicate each respective *S. cerevisiae* chromosome from where tRNA genes originate and arrows indicate the order of the tRNA arrays in the final tRNA neochromosome. **(B)** The synthetic bidirectional termination sites are based on the native rDNA replication fork blocking sites mediated by Fob1, harvested from various *Saccharomyces* species rDNA units. In natural replication fork blocking sites, DNA replication may only proceed in one direction (green arrow) and is blocked in the other direction (red line). In contrast, synthetic termination sites described consist of two RFBs from different *Saccharomyces* species, DNA replication readily passes through the first barrier but is blocked by the second barrier. This design is intended to minimize potential collision events between the DNA replication and RNAPIII transcription machinery.

The design of the tRNA neochromosome was framed around a series of design specifications (design features are summarized in Box 1), largely intended to maximize its stability and ensure tRNA functionality. One of our primary design considerations was to limit homology with the host genome by incorporating DNA sequences that are largely orthogonal to the wild-type and synthetic yeast genome sequences. Overall, the majority (∼89%) of the tRNA neochromosome sequence originates from non-*S. cerevisiae* yeast species. These orthogonal regions include tDNA flanking sequences, three origins of replication and four novel synthetic bidirectional replication termination sites assembled *de novo* and based on natural *Saccharomyces spp. Fob1* recognition sequences.

The tRNA neochromosome was designed to maintain tRNA copy number and codon specificity of all wild-type tRNA genes of *S. cerevisiae*. Since ensuring tRNA gene function took priority over orthogonality, tDNAs were not altered (excepting introns, which were removed from all 59 intron-containing tDNAs). To minimize host genome homology, each tRNA cassette incorporated flanking sequences from the yeasts *Ashbya* (*Eremothecium*) *gossypii* or *Eremothecium cymbalariae*. We selected these yeast species due to their close evolutionary relationship to *S. cerevisiae* (Dietrich *et al*., 2004; Wendland and Walther, 2011). Furthermore, *A. gossypii* lacks retrotransposons (Dietrich *et al*., 2013), and so is not populated by the repetitive solo LTR elements that commonly reside upstream of *S. cerevisiae* tDNAs. Each 500 bp 5’ flanking region was chosen to ensure a consistent spacing between each highly-transcribed tRNA gene sequence (Fig. 1A). The 40 bp 3’ region was incorporated to ensure efficient tRNA transcriptional termination mediated by poly-thymidine residues that constitute the RNAPIII terminator (Braglia *et al*., 2005). Since tRNA genes are transcribed from an internal promoter, we did not envision a significant disruption of tRNA transcription. Each tDNA was assigned to flanking sequences (Tab. S3) using a bioinformatic pipeline that additionally removes or alters unwanted elements from flanking sequences (Fig. S1A). These elements included residual start codons, LTR repeats from *E. cymbalariae* and overlapping regions from adjacent tDNAs (Fig. S1B). Furthermore, a proof-of-principle experiment demonstrated that synthetic copies of single-copy, essential tRNA genes (*SUP61* (*tS(CGA)C*), *TRT2* (*tT(CGU)K*) and *TRR4* (*tR(CCG)L*)) were capable of successfully complementing deletion of their wild-type counterparts, with no significant effect of cell phenotype (Fig. S1C). Similarly, removing an intron from the synthetic copy of tS(CGA)C did not result in a phenotype, in agreement with a previous study (Hayashi *et al*., 2019), although a potential precursor accumulation and a single downstream deleterious synthetic genetic interaction was observed for *tS(CGA)C* (Zhao *et al*., 2022).

### Automation of tRNA flanking sequence assignment

The unique structure of the tRNA neochromosome necessitated a computer assisted design approach to complement the design process. The bioinformatics algorithm and associated scripts (Fig. S1A) were used to streamline the assignment of tRNA genes to respective flanking sequences recovered from *A. gossypii* and *E. cymbalariae*.

Sixteen tRNA arrays (Fig. 1A), consisting of concatenated tRNA cassettes, housed tRNA genes from each of the sixteen respective *S. cerevisiae* chromosomes were re-organized so that relative tRNA gene orientation was unidirectional for each array. The unidirectionality was deemed potentially important to assure that the directions of replication forks and RNAPIII were always congruent. Furthermore, tRNA genes in tRNA arrays were aggregated by chromosome in a modular manner to both facilitate the construction of the tRNA neochromosome itself, and additionally, complement the loss of tRNA genes on individual synthetic chromosomes as part of the Sc2.0 project (Zhao *et al*., 2022) and to assist in debugging of synthetic chromosomes. In terms of additional evidence for function of individual synthetic tRNAs constructed according to this overarching design scheme, we note that the *chrII* and *chrXII* tRNA arrays, respectively, have previously demonstrated synthetic chromosome complementation through reversal of global upregulation of the *synII* translational machinery (Shen *et al*., 2017b) as well as restoration of a fitness defect in *synXII* (Zhang *et al*., 2017).

### Neochromosome structure

The overall structure of the tRNA neochromosome (Fig. 1A) was designed to encompass the fundamental features of typical eukaryotic chromosomes, including a centromere, telomeres and origins of replication. The backbone for tRNA neochromosome assembly comprised the pRS413 centromeric vector, which itself houses the centromere region of chromosome VI (*CEN6*) fused to an origin of replication (*ARSH4*) and a *HIS3* marker (Sikorski and Hieter, 1989). Three orthogonal origins of replication from the yeast *Candida glabrata* (Tab. S7), were introduced and positioned so as to ensure relatively even tRNA neochromosome replication (Fig. 1A) with ‘early’ and ‘strong’ replication origins preferentially selected. Our circular design also enables us to chemically extract the neochromosome and transform it into a new yeast background later (Noskov *et al*., 2011), and also to conditionally linearize by introducing functional telomeres in any defined region using the telomerator system (Mitchell and Boeke, 2014) (*cf*. Fig. 3A).

### Minimizing replication stress

A key consideration in our design was to maximize tRNA neochromosome stability, which included an overall aim of minimizing replication stress caused by polar replication fork collisions with tRNA gene transcription. tRNA genes are among the most highly transcribed in the cell (Dieci *et al*., 2007; Waldron and Lacroute, 1975). It is also known that replication fork stall events in eukaryotes are influenced by the orientation of tDNA transcription relative to replication. When oriented to oppose incoming replication forks, they can induce blockage of the latter and induce chromosomal instability (Deshpande and Newlon, 1992; Hamperl *et al*., 2017). Therefore, to reduce replication stress and DNA damage caused by head-on replication fork stall events, we oriented tRNA arrays so that the direction of both replication and tRNA transcription were concordant (Fig. 1). These formed mega-arrays which were designed to be of similar size (approx. 20 kb each), to ensure replication progression as evenly as possible across the resulting ∼40 kb replicons.

An inevitable challenge caused by the presence of multiple replication origins in our design is that a replication fork may progress beyond regions carrying co-oriented transcription units and subsequently collide with tRNA transcription in opposing orientation. To limit this issue and minimize potentially destabilizing effects of head-on replication fork collisions with components associated with active tRNA transcription, we introduced four cassettes designed to block DNA replication in both directions. Our strategy intended to emulate the protective role of replication termination sites of the rDNA locus (Brewer *et al*., 1992), which are known to prevent head-on collisions between the replication fork and the RNA Polymerase I (RNAPI) transcriptional machinery of rDNA (Rothstein *et al*., 2000; Takeuchi *et al*., 2003). Each of the four designer bidirectional replication termination sites (*TERI to TERIV;* (Tab. S7) were designed to house two conserved Fob1 recognition sites, in opposite direction, with sequences based on the rDNA sequences of four non-*S. cerevisiae Saccharomyces* species (Fig. 1B).

### An orthogonal recombinase system for tRNA copy number adjustment

The Sc2.0 project leverages SCRaMbLE (Synthetic Chromosome Rearrangement and Modification by *LoxP*-mediated Evolution) to generate combinatorial diversity in strains housing synthetic chromosomes (Shen *et al*., 2016). To explore questions of tRNA genetics such as minimal and/or optimal combination of tRNA genes needed under a given condition, we introduced a second, independent site-specific recombinase system, based on the Dre-*rox* system (Liu *et al*., 2018a; Sauer and McDermott, 2004), to facilitate future studies. The positioning of *rox* recombination sites both upstream and downstream of each tRNA cassette (Fig. 1A) is intended to provide the means to stochastically alter tDNA copy number. There is no evidence of any cross-reactivity between the Cre-*lox* and Dre-*rox* recombination systems (Anastassiadis *et al*., 2009). A preliminary proof-of-principle experiment demonstrated that the Dre-*rox* system successfully induced a reduction in the number of tRNA cassettes on a tRNA array (Fig. S1D). It should be noted that the tRNA neochromosome was constructed in a wild-type background with all native tRNA genes present. In the future, we will adjust tRNA copy number using the Dre-*rox* system once the fully synthetic Sc2.0 strain is available.

### Neochromosome construction

To construct the tRNA neochromosome from its constituent tRNA arrays, we employed an approach similar to the SwAP-In method described previously, but without using a natural template (Annaluru *et al*., 2014; Richardson *et al*., 2017). This evolved into the eSwAP-In approach described previously (Mitchell *et al*., 2021) and is referred to as such henceforth. This process applied recursive rounds of *in vivo* homologous recombination between a circular, intermediate “acceptor” assembly and a single “donor” tRNA array. Integration of each tRNA array was facilitated by two regions of homology: the outermost ∼400 bp to ∼600 bp DNA sequence already present in the growing neochromosome, and a 500 bp “Universal Homologous Region” (UHR) consisting of randomly-generated DNA. The repeated swapping of the *LEU2* and *URA3* selective markers facilitated selection for each round of integration (Fig. 2A). Successful integration of each tRNA array was verified through PCR using specific primers designed to anneal with the 3’ and 5’ flanking region of each tRNA cassette (Fig. S2D, Tab. S4). The tRNA neochromosome was constructed in two independent strain backgrounds, in parallel, through a series of 15 subsequent rounds of integration into a circular acceptor vector housing the *chrVI* tRNA array. The first of these strains was the wild-type BY4741, the second strain housed synthetic chromosome III, VI and the right arm of IX (*synIII/VI/IXR*) (Richardson *et al*., 2017). Subsequently, the circular tRNA neochromosome constructed in *synIII/VI/IXR* was subjected to chemical extraction and spheroplast transformation into a ‘fresh’ BY4741 strain resulting in strain YCy1576.

**Fig. 2.**
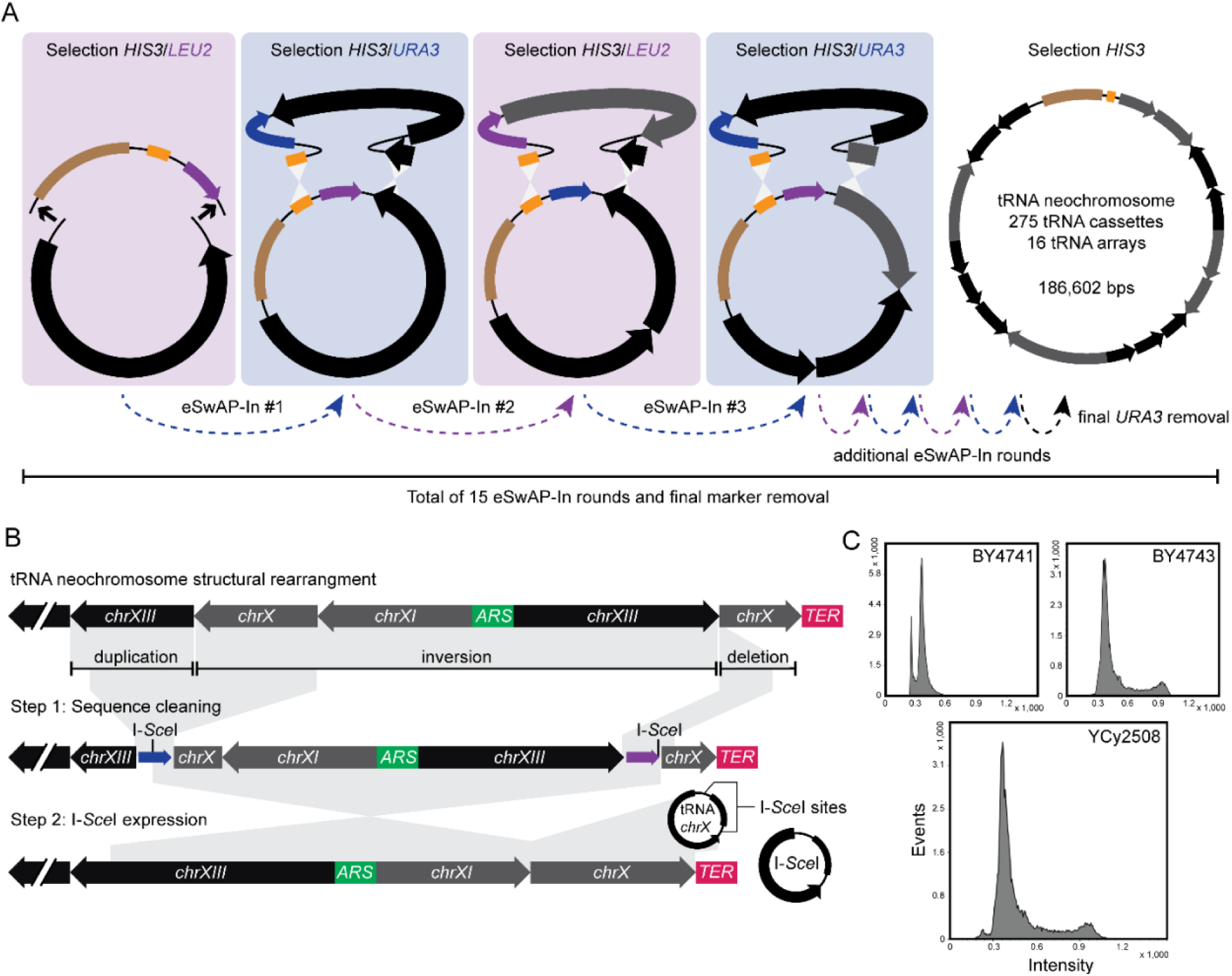
Construction, debugging and discovery of a whole genome duplication. **(A)** tRNA neochromosome construction was performed in a stepwise manner through repeated rounds of *in vivo* homologous recombination in yeast. An exception was the first tRNA array (*chrVI*), which was instead ligated into a *LEU2* eSwAP-In vector (indicated by small arrows in the first panel) to provide a start point for tRNA neochromosome construction and the *CEN*/*ARS* and *HIS3* marker required for *in vivo* function. The brown bar denotes the pRS413 region of the eSwAP-In vector, black and gray arrows denote tRNA arrays. *In vivo* homologous recombination is facilitated by two homology arms incorporating the last 350-680 bp sequence from the previous array and the 500 bp ‘universal homologous region (UHR; shown in orange). Selection is conferred by the recursive swapping of *LEU2* and *URA3* markers. Following the final round of tRNA array integration, the *URA3* marker was removed to facilitate tRNA neochromosome linearization. **(B)** Following tRNA neochromosome construction, a complex structural rearrangement was observed consisting of a duplication, inversion and deletion. The debugging process was performed in two steps; to first remove the duplicated sequence and the majority of tRNA array *chrX* via the integration of a *URA3* and *LEU2* marker coupled to I-*Sce*I recognition sites. In the second step I-*Sce*I was overexpressed in the resulting strain supplied with a plasmid containing a *chrX* tRNA array flanked by I-*Sce*I recognition sites. Induction of I-*Sce*I results in two cuts in the tRNA neochromosome to repair the inversion as well as the release of tRNA array *chrX* from the plasmid to fill the gap resulting in the tRNA neochromosome that met our basal design. **(C)** The tRNA neochromosome strain showed an unexpected whole genome duplication and a *MAT***a**/**a** genotype which were verified by flow cytometry, mating-type tests (Fig. S3) and diagnostic PCRs (data not shown).

### Neochromosome linearization and repair of structural variations

Initial tRNA neochromosome linearization experiments unexpectedly revealed structural variations of the tRNA neochromosome constructed in both strain backgrounds (see below, and Fig. S2C). Notably, pulsed-field gel electrophoresis (PFGE) and next-generation sequencing revealed complex structural variations of varying genomic sizes, including a doubling in size, of one BY4741 tRNA neochromosome variant (Fig. S2C). In contrast, whole genome sequencing of the tRNA neochromosome constructed in *synIII/VI/IXR* (YCy1576) revealed a single complex structural variation, consisting of an inversion, duplication and deletion (Fig. 2B). Furthermore, a total of eleven minor nucleotide variations outside the pRS413 backbone were detected, with nine having no predicted impact and two potentially inactivating one *rox* site each (Tab. S6).

We elected to repair the structural variations of YCy1576 to generate the circular tRNA neochromosome consistent with our basal design. The structural variations of the resulting strain, YCy1677, were subsequently repaired by employing a two-step procedure mediated by the I-*Sce*I homing endonuclease to remove the duplicated region, introduce the missing tRNA cassettes and repair the inversion (Fig. 2B). The final NGS validated strain (YCy2508), was used for subsequent experiments.

### Spontaneous doubling in yeast genome ploidy and tRNA neochromosome instability in haploid cells

Unexpectedly, we observed a uniform increase in host strain ploidy from a haploid to a homozygous diploid (*i.e.* from 1*n* to 2*n* BY4741 *MAT***a**/**a** genotype). This was confirmed via the strain’s ability to mate with *MATα* strains, PCR-based mating type tests (Huxley *et al*., 1990) and flow-cytometry quantification of DNA content (Fig. 2C and Fig. S3). In addition, flow cytometry revealed the same doubling in DNA content in a separate strain housing a tRNA neochromosome which had *not* undergone spheroplast transformation (Fig. S4). Spontaneous alterations of cell ploidy in yeast have been associated with a stress response (Harari *et al*., 2018b). Therefore, we hypothesized that the increase in cell ploidy was a compensatory response to the presence of the tRNA neochromosome in wild-type haploid cells.

Given the apparent selective force manifesting as an increase cell ploidy, we later elected to maintain the tRNA neochromosome in the *MAT***a**/**a** genome-duplicated BY4741 background (YCy2508) for all subsequent functional studies. This rationale was supported by time-course experiments performed in independent, tRNA neochromosome-transplanted haploid strains (BY4741 and BY4742), which revealed an apparent increase in tRNA neochromosome stability in the diploid background. Following haploid validation *via* flow-cytometry (Fig. S5), an increased number of internal deletions were observed in independent, freshly-transformed haploid strains ((BY4741 (YCy2837), BY4742 (YCy2838)) compared to that of the *MAT***a**/**a** diploid (YCy2508) and *MAT***a**/*α* diploid (YCy2844) strains. After approximately 50 generations, deletions were detected in two out of eight diploid isolates compared to 7 out of 8 haploid isolates (Fig. S5).

### Linearization and sequencing of tRNA neochromosome strains reveals extensive genomic plasticity

To probe the effect of linearizing the tRNA neochromosome at different locations, a total of nine locations were selected (generating strains YCy2670 to YCy2678). Four were linearized at *TER* sites, two close to the centromere and three at origins of replications (*ARSI* to *ARSIII*) (Fig. 3A). Successful linearization and tRNA neochromosome size were verified via PFGE analysis. With the exception of one mitochondrially-deficient rho^-^ mutant (YCy2677), we observed no significant growth difference between isolates housing either circular or linearized variants (Fig. 3B and S6). Notably, with the exception of potential aneuploidies and sequence variations in several independent strains linearized at non-*TER* positions (Fig. S7), disruption of the replication profile of the tRNA neochromosome through linearization at origins of replication produced no significant impact on cell phenotype compared to replication termination sites (*TERI-IV*) under 22 growth conditions (Fig. S6). Overall, our results suggest that there is no significant difference in cell growth rate between linear or circular tRNA neochromosome variants, and that the location chosen for linearization has no major impact on cell growth.

**Fig. 3.**
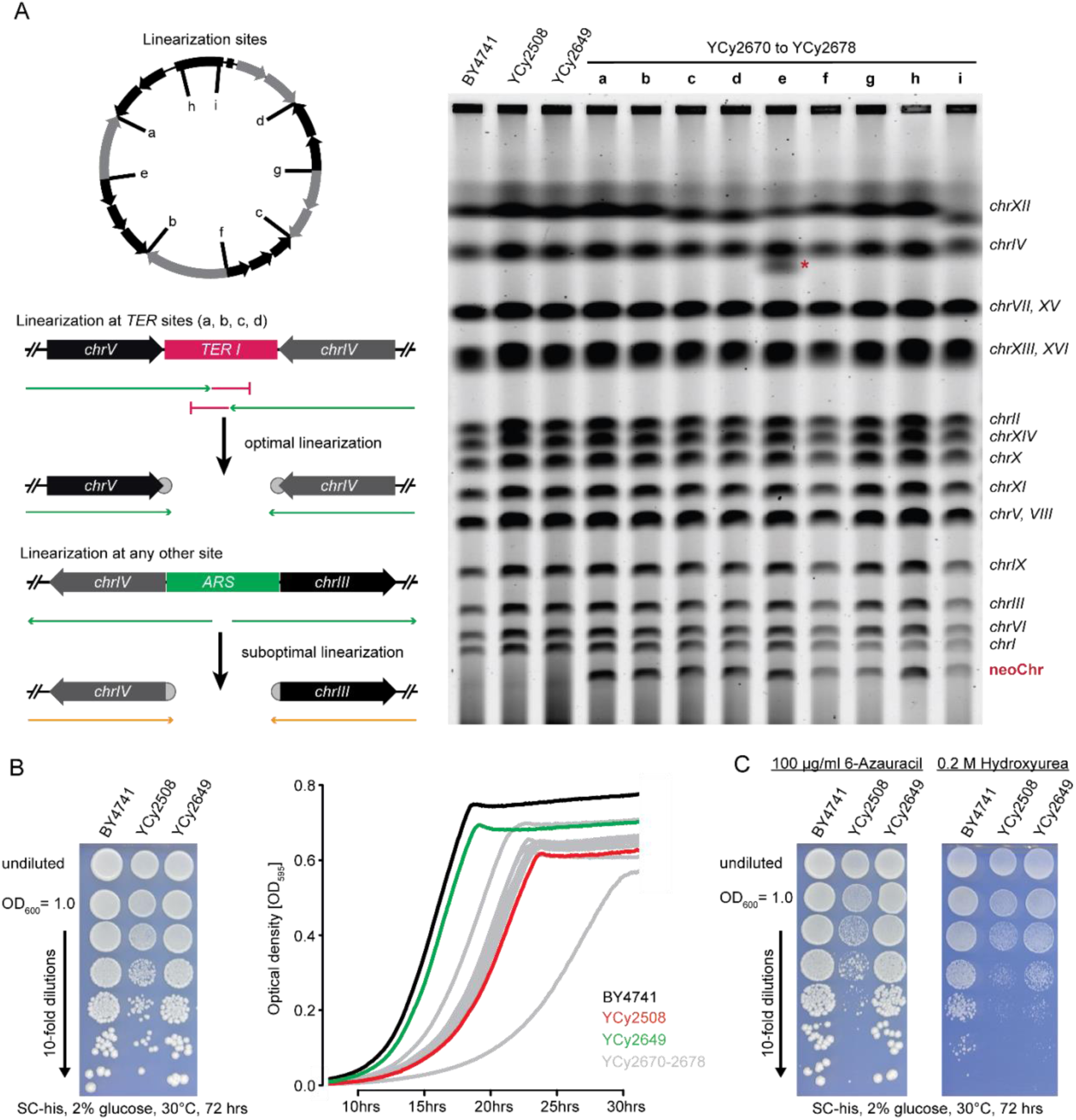
Linearization and phenotypic characterization of the tRNA neochromosome. **(A)** The panel on the left indicates linearization at sites predicted to be optimal for the proper alignment of DNA replication and *RNAPIII* transcription (*i.e.* at *TER* sites), or suboptimal for tRNA neochromosome linearization (*i.e.* at origins of replication). DNA replication aligned with transcription is indicated by green, DNA replication blocking by red and colliding DNA replication and transcription by orange arrows. The panel on the right shows pulsed field gel electrophoresis (PFGE) of tRNA neochromosome strains and respective controls. The circular tRNA neochromosome (YCy2508) cannot be visualized by PFGE analysis due to topological trapping in agarose fibres. However, the linearized versions (YCy2670 to YCy2678) show a band on the expected height (indicated by neoChr). All chromosome bands are as expected, except for strain YCy2674 where a fraction of *chrXII* (red asterisk) shows a partial loss of the rDNA repeat (supported by NGS data, data not shown). **(B)** Phenotypic growth assays indicate that the tRNA neochromosome strains have a growth defect which is recovered upon loss of the tRNA neochromosome. The growth differences between circular and linear strains are marginal (with the exception of one strain which was later found to have partial loss of mitochondrial DNA). **(C)** The tRNA neochromosome strains were benchmarked under 22 separate media conditions (*cf*. Fig. S6) and compared to BY4741 + pRS413 and YCy2649. Only in the presence of 6-Azauracil or 0.2 M Hydroxyurea the phenotype becomes more severe indicating transcriptional and DNA-replication stress, respectively.

### tRNA sequencing to characterize tRNA neochromosome expression

To examine whether tRNA genes on the tRNA neochromosome are transcribed when removed from their native genomic context, we performed tRNA sequencing (tRNAseq). Due to extensive tRNA secondary and tertiary structures combined with the indistinguishability of individual locus expression of multigene tRNA species after tRNA processing, traditional RNA sequencing approaches are limited in terms of unbiased and global tRNA characterization. To overcome these challenges, we implemented a dedicated workflow and developed sensitive computational approaches for transcript-level quantification to distinguish closely related tRNA species (Fig. 4A). We performed tRNAseq on four strains with three biological replicates each, including the haploid and homozygous diploid controls (YCy2409, YCy2649), the circular (YCy2508) and a linear tRNA neochromosome strain (YCy2671). Leveraging leading and tailing pre-tRNAs sequences we were able to identify the genomic origin of 255 of the 275 tRNAs based on their pre-tRNAs and estimate their fractional expression (Fig. 4A and 4B, Supplementary Data S2).

**Fig. 4.**
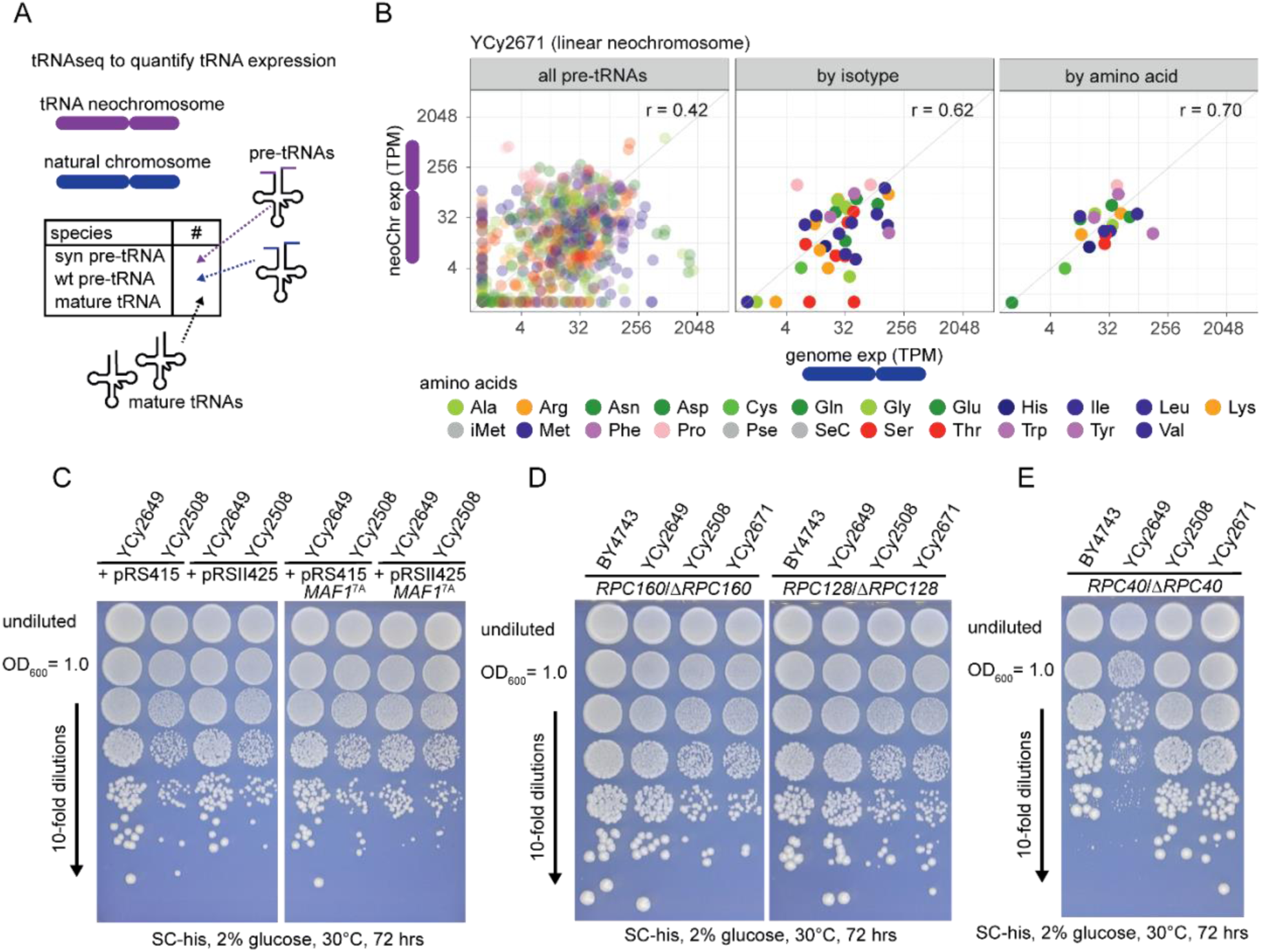
tRNA expression analysis and phenotypic impact of RNAPIII activity reduction. **(A)** tRNAseq enables differentiation between the different tRNA loci based on the fraction of pre-tRNAs. Leading, tailing and intron sequences allow a clear separation of transcript origin. **(B)** tRNAseq data shows that the expression levels of individual synthetic and native tRNA genes varies, sometimes dramatically. However, sorting tRNA expression by isotype or by amino acid indicates that overall expression levels appear to be relatively well-balanced whether the source gene is native or synthetic. This is consistent with a general tRNA expression buffering mechanism within yeast cells. **(C)** Overexpressing the constitutively active MAF1^7A^ allele rescues the observed phenotype in the tRNA neochromosome strain. However, the growth of the control strain is reduced and almost matches the growth defect of the tRNA neochromosome strain when transformed with a high copy plasmid (pRSII425) overexpressing MAF1^7A^. Interestingly, the growth of the tRNA neochromosome strain is not further reduced, potentially indicating that Maf1 is fully active in this strain to compensate the consequences of tRNA overabundance. **(D)** Reducing RNAPIII activity by deleting single subunit alleles in the homodiploid did not show strong effects on growth recovery, neither in the circular nor in a linear tRNA neochromosome strain. *RPC160* and *RPC128* are shown as examples in comparison to the homodiploid control. **(E)** *RPC40,* in contrast, produced a recovery in growth phenotype for both the circular and linear tRNA neochromosome strain. The control strain (BY4743 + pRS413) shows two types of colony sizes. Sequencing of two big and two small clones by nanopore sequencing revealed the small clones have a low abundance of mitochondrial DNA in comparison to the large clones (data not shown).

Our analysis of pre-tRNAs revealed a similar expression pattern between the detected pre-tRNA abundances from the tRNA neochromosome and the genome aggregated by the 20 amino acids or 42 isoacceptor types, although individual tDNA expression was not so well correlated (Fig. 4B). Furthermore, our results revealed no systematic alteration of tRNA expression caused by the removal of introns (Fig. S8).

Northern blot analysis previously indicated a potential 5’ tRNA maturation deficiency for the single-copy synthetic tRNA_Ser_^CGA^ (syn*SUP61*) (Zhao *et al*., 2022). Global tRNA expression analysis (Fig. S8 and S9) did not provide any evidence for accumulation of pre-tRNAs, which would provide evidence for systematic tRNA maturation deficiencies on genes encoded by the tRNA neochromosome. Indeed, the tRNA_Ser_^CGA^ pre-tRNA was one of the few isoacceptors that were not detected following global tRNAseq after data normalization (Fig. 4B and S8, Supplementary Data S2), and so we cannot definitively determine if any of the other 20 “missing” tRNA genes may be affected.

### Phenotypic characterization of the tRNA neochromosome

We observed a growth impact in homozygous diploid cells housing both linearized and circular variants of the tRNA neochromosome (YCy2670 to YCy2678), with an extended lag phase and an increase in doubling time of 19.2% (92.3 +/- 2.7 min vs 110.0 +/- 2.6 min; a rho^-^ mutant was excluded from this calculation) compared to haploid and homozygous wild-type cells (Fig. 3B). To confirm that the tRNA neochromosome is the cause of the growth defect, we selected for its spontaneous loss on nonselective media. The resulting isolate (YCy2649) reverted to a normal phenotype, demonstrating that the tRNA neochromosome itself was responsible.

To further assess the potential cause of the phenotype, we subjected strains housing circular and linear versions of the tRNA neochromosome to a total of 22 solid media growth conditions (Fig. S6). We observed a relative growth defect compared to the control under all conditions, with a more significant defect in the presence of 6-azauracil and hydroxyurea, indicating transcriptional elongation stress and replication stress, respectively (Fig. 3C and S6). The latter phenotype on hydroxyurea also revealed a similar growth defect on the *MAT***a**/**a** diploid control strain (absent of the tRNA neochromosome), suggesting a potential phenotype caused by a change in the host genome, or through whole genome duplication. Notably, a similar effect for hydroxyurea was observed for growth phenotypes of a yeast strain possessing a *synVII* aneuploidy (Shen *et al*., 2022). We observed no difference between the circular and linear variations of the tRNA neochromosome (Fig. S6).

Given the relative doubling in the number of cellular tRNA genes, we set out to explore if reducing tRNA transcription might alleviate the observed cellular burden. RNAPIII is negatively regulated in a global manner by Maf1 (Pluta *et al*., 2001). To increase gene copy number, two additional *MAF1* variants were introduced onto a high-copy (2-micron) plasmid under control of their natural promoter: the wild-type variant and a previously described constitutively-active *MAF1^7A^* variant (Huber *et al*., 2009). Both were introduced into YCy2649 and YCy2508. Furthermore, a second series of experiments investigated a reduction in RNAPIII abundance by individually deleting one copy of all ten essential RNAPIII specific subunits (Wild and Cramer, 2012) of the seventeen-protein core complex individually, resulting in ten heterozygous *MAT***a**/**a** strains.

Overall, our results show that neither *MAF1* variant, nor the majority of RNAPIII deletions, produced any significant phenotypic effect on strains housing the tRNA neochromosome (Fig. 4 C-D and S10). However, an exception may be observed following a slight phenotype recovery in a strain containing an *RPC31* deletion. Rpc31 is notable due to its apparent role in regulating RNAPIII activity (Huet *et al*., 1985), and mutation of *RPC31* can suppress a Δ*MAF1* phenotype (Cieśla and Boguta, 2008; Thuillier *et al*., 1995).

We further decided to generate heterozygous deletions of the two large subunits of RNAPI (*RPC128* and *RPC160*) and two shared RNAPI and RNAPIII subunits (*RPC19* and *RPC40*). Our results (Fig. 4 E) indicate that eliminating one copy of *RPC40* deletion produced an apparent recovery of growth for tRNA neochromosome strains (although large and small colony sizes for the control strains were notable, with Nanopore sequencing revealing a reduction in mitochondrial DNA for the smaller colonies). No significant changes in phenotype were observed following a reduction in *RPC128*, *RPC160* and *RPC19* abundance. Rpc40 is necessary for the assembly of RNAPI and RNAPIII (but not RNAPII). Moreover, Rpc40 is necessary for retrotransposon insertion specificity (Bridier-Nahmias *et al*., 2015; Lalo *et al*., 1993; Mann *et al*., 1987). Taken together, our results suggest that increased abundance of tRNA genes may at least be partially responsible for the observed phenotype. The combined reduction of both RNAPI and RNAPIII abundance may be beneficial to recover the growth phenotype.

### Transcriptome and proteome analysis of tRNA neochromosome strains

Transcriptomics and proteomics were later performed to further investigate the effect of the tRNA neochromosome. RNA sequencing (RNA-Seq) was performed on the circular and seven linear tRNA neochromosome variants, as well as the haploid and homozygous diploid control strains (Fig. 5A and Fig. S11), respectively. Our results show a general upregulation of amino acid biosynthesis-related transcripts, and a downregulation of ribosomal protein transcripts for all strains housing the tRNA neochromosome (Fig. 5A). Proteomics was performed on the circular (YCy2508) and nine linear tRNA neochromosome variants (YCy2670-2676) (Fig. 5B and Fig. S12), as well as the haploid (BY4741), homozygous diploid (YCy2649) and diploid (BY4743) control strain, revealing no significant changes at the individual protein level. However, at the collective level we observed an increase in proteins associated with amino-acid biosynthesis and a reduction in proteins associated with the ribosome (Fig. 5B), correlating well with our transcriptomic data. Overall, neither transcriptome nor proteome analysis indicate a strong differential regulation of the DNA damage response.

**Fig. 5.**
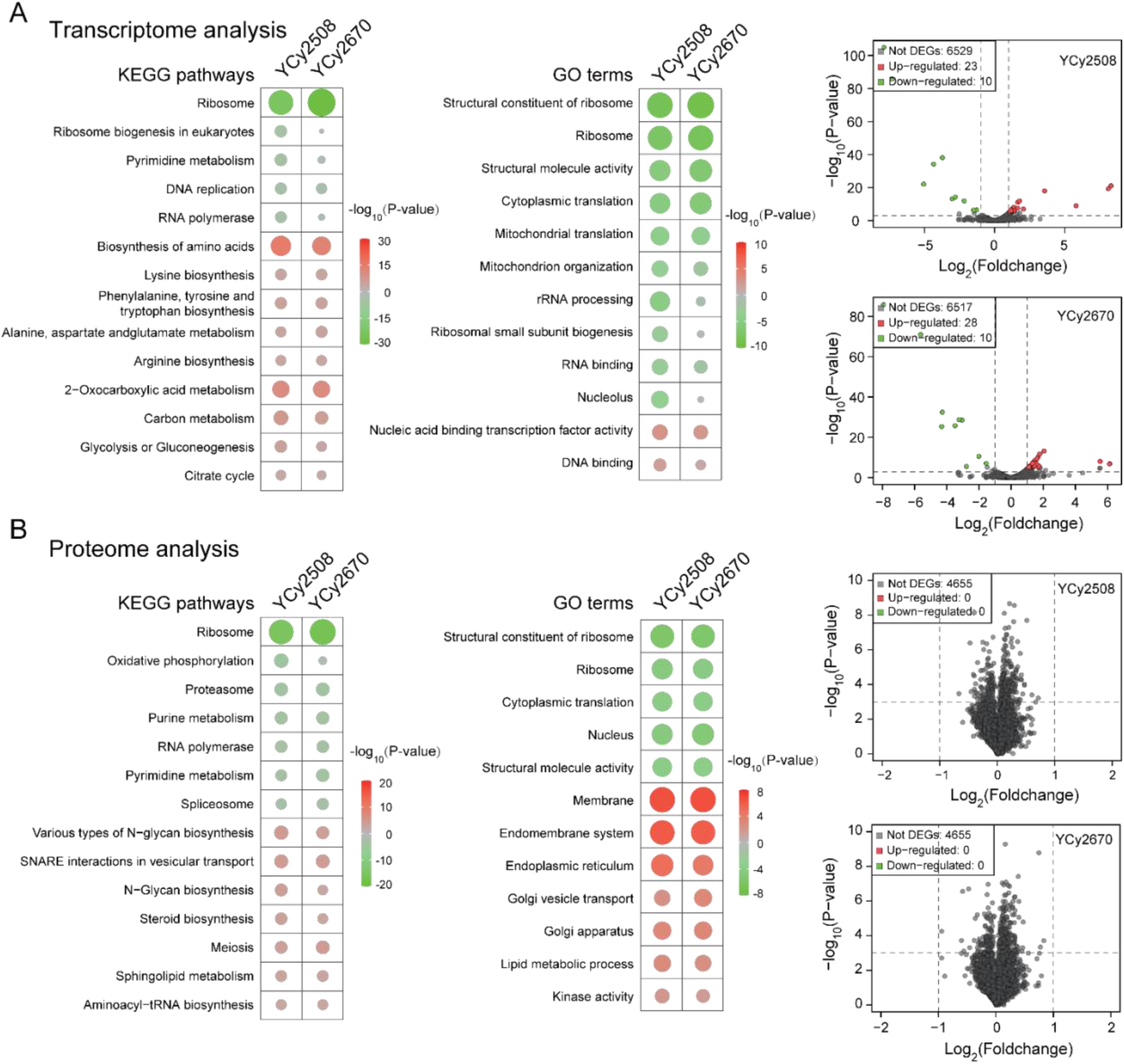
Transcriptome and proteome analyses indicate a general upregulation of amino acid biosynthesis and downregulation of ribosomal proteins. **(A)** Differential expression of circular (YCy2508) and linear (YCy2670) tRNA neochromosome strains normalized to the homozygous diploid control strain (YCy2649). KEGG (left panel) and GO (right panel) analysis demonstrate a general upregulation of amino acid biosynthesis and a down regulation of translational processes. The volcano plots indicate that 33 and 38 individual mRNA transcripts associated with the latter are upregulated or downregulated, respectively. Up-regulated and down-regulated features are labeled in red and green, respectively. **(B)** Proteomic analysis of the circular and linear tRNA neochromosome strains (YCy2508 and YCy2670), respectively, normalized to the homozygous diploid control strain (YCy2649). KEGG and GO analysis globally show a similar pattern to that of transcriptomics. However, an additional increase in membrane-associated proteins and endoplasmic reticulum can be observed. Up-regulated and down-regulated features are labeled in red and green, respectively.

### Nucleosome mapping

We subsequently investigated whether the clustering of tDNAs on the tRNA neochromosome might affect the general organization of its chromatin structure. Nucleosome occupancy maps were generated to investigate any potential for aberrant chromatin structure, such as mispositioning or nucleosome absence. Our results show that nucleosome positioning for genomic tRNAs are similar in all strains tested (Fig. 6A). Notably, with the exception of a slight apparent variance in linker length, nucleosome positioning on both tRNA neochromosome variants is similar to that of genomic tRNAs, suggesting a relatively normal chromatin structure. Our data suggests that the typical 20-bps linker between two nucleosomes is reduced in the tRNA neochromosome variants, presumably due to the tight clustering of tDNAs. Our observations suggested that active transcription of tRNA genes may contribute to upstream and downstream nucleosome positioning. Aside from the high density of neochromosome tDNAs and an apparent difference in linker length, nucleosome positioning adopts a rather orthodox pattern and is nearly indistinguishable from native positioning.

**Fig. 6.**
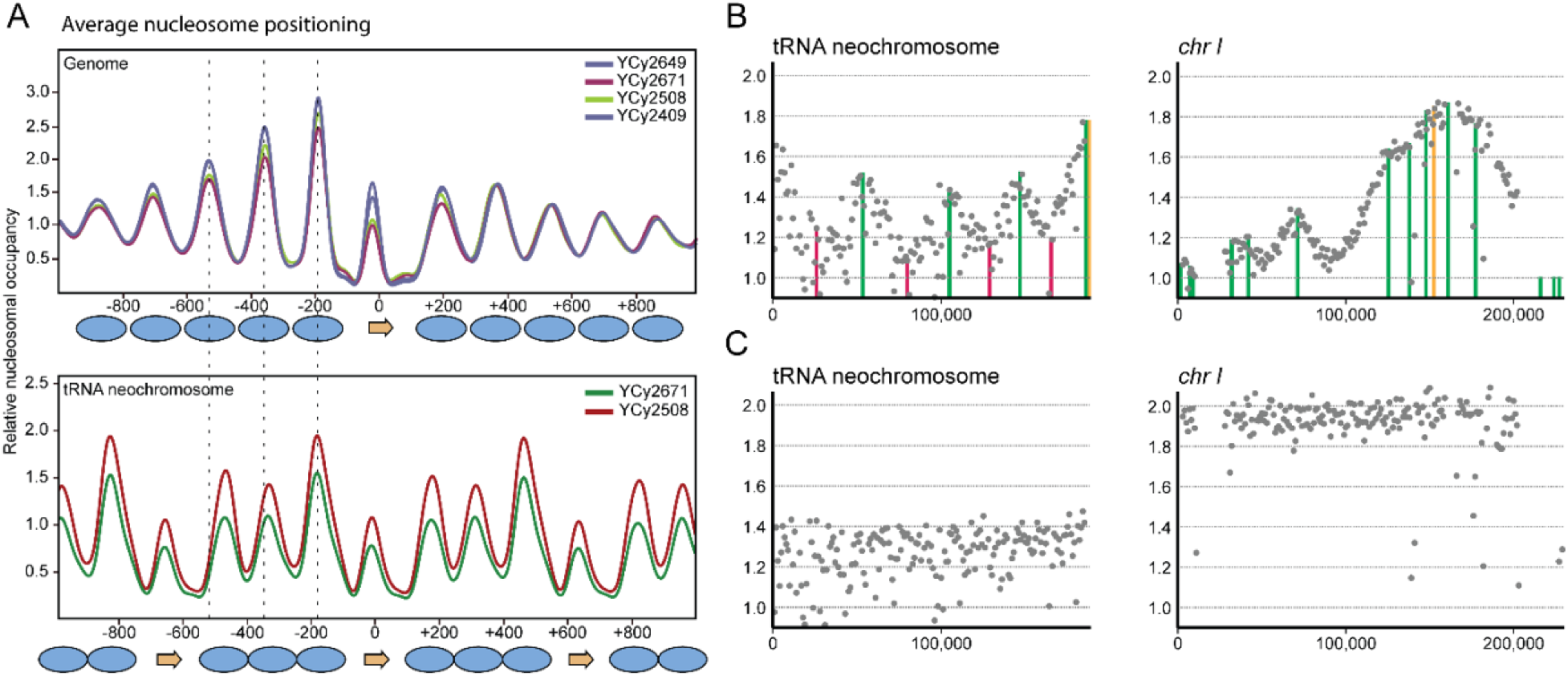
Nucleosome positioning and replication pattern of the tRNA neochromosome. **(A)** Averaged nucleosome positioning around tRNA genes on the native chromosomes are indistinguishable within all tested strains (top panel). The tRNA neochromosome nucleosome positioning is similar (bottom panel). However, based on the high density and the distance between two tRNA genes, the nucleosomes appear more compacted, as indicated by the dotted lines aligned to the dyad position of the three nucleosomes upstream from the tRNA in the genome (top panel). **(B)** Replication pattern analysis indicate the functionality of origins of replication (green lines) and their time-point of activation within the cell cycle for the circular tRNA neochromosome. Peaks correspond to initiation events and valleys correspond to termination of DNA replication. In case of the tRNA neochromosome, all origins from *C. glabrata* show apparent initiation events and associated valleys corresponding to termination sites (red lines). The position of the centromere is visualized by an orange line. **(C)** Marker frequency analysis of cells synchronized to G1 show a lower abundance of the tRNA neochromosome compared to the native chromosomes. Synchronization at the end of S-phase revealed a similar relative abundance between the tRNA neochromosome and native complement (*cf*. Fig. S13).

### Deep-sequencing to determine the tRNA neochromosome replication profile

The tRNA neochromosome houses four origins of replication and four bidirectional replication termination sites. To explore how the tRNA neochromosome replicates, we performed deep sequencing of replicating cells normalized to the sequencing reads of non-replicating cells in both a circular (YCy2508) and a linear (YCy2671) tRNA neochromosome variant (Fig. S13). Our results show maxima of sequencing coverage co-localizing with each origin of replication, and minima associated with the designed bidirectional replication termination sites (Fig. 6B). Within the analysis it was observed that the tRNA neochromosome tends to have lower sequencing coverage compared to the native chromosomes. To further investigate this observation, cells were normalized to G1 (α factor arrest) (Fig. 6C and Fig. S13). Interestingly, the tRNA neochromosome displayed an approximate abundance of 0.8 relative to the other chromosomes in G1 phase, suggesting a heterogeneity in tRNA neochromosome copy number or partial tRNA neochromosome loss within the cell population. These data suggest that the tRNA neochromosome has the same abundance as the native chromosomes before cell division, but potentially aberrant segregation leads to the observed G1 effect, which we speculate may be one potential cause of the observed reduced growth rate.

### Intranuclear localization of the tRNA neochromosome

Evidence suggests that tDNAs influence the overall organization of the genome in the nuclear space (Chen and Gartenberg, 2014; Duan *et al*., 2010; Thompson *et al*., 2003). Recently, the investigation of a yeast strain devoid of tRNA genes on *chrIII* and other synthetic chromosomes showed an impact on centromere clustering, but no significant influence on the large-scale architecture of the genome (Hamdani *et al*., 2019; Mercy *et al*., 2017). Given the designer structure of the tRNA neochromosome, we explored its position within the nuclear space and whether it modifies the global genome architecture using a combination of fluorescence *in situ* hybridization (FISH) and capture of chromosome conformation (Hi-C; Lieberman-Aiden *et al*. (2009)).

FISH experiments were performed using separate probes for *rox* recombination sites and rDNA repeats (5’Cy3 and 5’FAM, respectively), to identify the position of the tRNA neochromosome in the nuclear space. Our results show that the tRNA neochromosome does not co-localize with the rDNA signal in most cells of the homozygous diploid strain (YCy2508), with approximately half of the cells showing a clear separation of the two regions (Fig. 7A). We validated the FISH results using live-cell imaging (Fig. 7B). A Transcription Activator-Like Effector construct (TAL-GFP) was designed to target *rox* sites and the nucleolus marked with Nop10-mCherry. We observed foci in cells housing the tRNA neochromosome, again with no significant co-localization of the *rox* signals and the nucleolus (Fig. 7A and 7B). Overall, FISH and co-localization studies demonstrate no obvious co-localization of the tRNA neochromosome with the nucleolus.

**Fig. 7.**
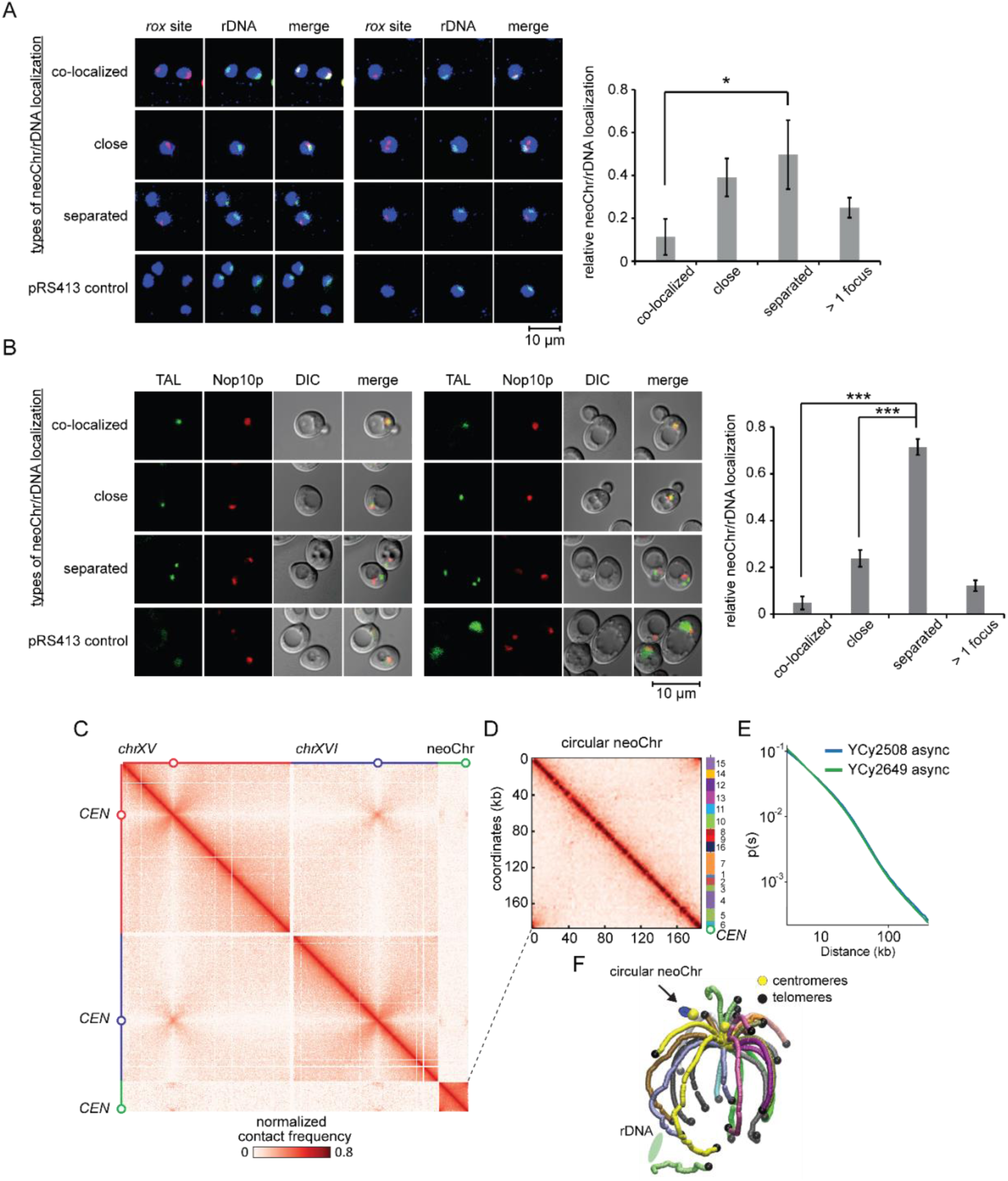
Intracellular localization of the tRNA neochromosome. **(A, B, F)** The tRNA neochromosome shows no obvious colocalization with the nucleolus. **(A)** The left panel displays representative FISH results visualizing the observed modes of tRNA neochromosome localization (red) relative to the rDNA locus (green) in strain YCy2508. The bottom panel on the left represents a pRS413 control absent of *rox* localization. Percentage of the different localization modes are given in the histogram on the right (three strains n = >100, respectively). * = 0.01<P<0.05. **(B)** Imaging results of live cells (JSY396 and JSY397). TAL: *rox*-probe-GFP, Nop10p:mCherry. The percentage of the different localization modes are given in the histogram. The percentage of the different localization modes are given in the histogram (three clones n = >250, respectively). *** = P<0.0001. **(C)** Magnification of the genome-wide contact map of the circular tRNA neochromosome strain. The x and y-axis represent chromosomes XV, XVI and the tRNA neochromosome, respectively. The color scale reflects the frequency of contacts between two regions of the genome (arbitrary units) from white (rare contacts) to dark red (frequent contacts). **(D)** Separate visualization of the circular tRNA neochromosome (YCy2508) showing the expected circularization signal at the edges of the contact map. No rearrangement is present. **(E)** Plot of the contact probability p as a function of genomic distance s (log scale) for the contact maps of either the tRNA neochromosome strain (YCy2508) or a strain (YCy2649), during exponential growth. **(F)** 3D representation of the contact map showing the clustering of the tRNA neochromosome (blue chromosome) in the vicinity of the centromere cluster and distal to the rDNA locus.

We performed Hi-C of circular (YCy2508) and linear tRNA neochromosome variants (YCy2670-YCy2672). Furthermore, Hi-C is a convenient approach to investigate structural genomic rearrangements in genomes (Marie-Nelly *et al*., 2014). Here, the contact maps of both variants displayed a smooth pattern, with no evidence of unwanted structural rearrangement, further validating tRNA neochromosome integrity (Fig. 7D). In addition, no obvious difference was observed between the circular and linear variants (data not shown). Intrachromosomal contacts were slightly enriched compared to native wild type yeast chromosomes. The greater intrinsic contacts observed within the tRNA neochromosome itself could correspond to a higher RNAPIII protein occupancy and transcription levels (Bignaud *et al*., 2022). In addition, the contact maps show that the tRNA neochromosome centromere is enriched in contacts with the native centromeres (Fig. 7C and 7F). The 3D representation of the 2D contact maps using ShreK3D (Lesne et al., 2014) was reminiscent to those obtained for diploid wild type (BY4743). The plot of the contact frequency as a function to the genomic distance computed over the entire genome (p(s) curve; materials and methods) was also conserved between the genomes of the tRNA neochromosome strain and a wild type diploid strain, suggesting that the presence of the tRNA neochromosome has no direct impact on the average conformation of the native genome (Fig. 7E).

## Discussion

We have described here the design, construction and characterization of a tRNA neochromosome, a fully synthetic designer chromosome which exists as an additional copy to the sixteen native chromosomes of *S. cerevisiae*. The design of the tRNA neochromosome was custom for the purposes of the Sc2.0 project and represents the first Sc2.0 synthetic chromosome designed without a natural template. This design was influenced heavily by engineering concepts common to synthetic biology, such as hierarchy, modularity and orthogonality, which helped to reconcile its structural complexity and were intended to maximize its stability *in vivo*.

We observed a remarkable plasticity of both the tRNA neochromosome and the host genome following construction. We speculate that the spontaneous increase in cell ploidy is a compensatory response to a cellular burden caused by a relative doubling in tRNA genes and consequent tRNA overabundance in a wild-type cell. The total RNA mass within a yeast cell consists to 80-90% of rRNAs, 10-15% of tRNAs, 3-7% of mRNAs and a small fraction of other RNAs (*i.e*. snRNAs, lncRNAs etc). However, if RNA mass is transferred to the actual number of molecules, tRNAs are the most abundant RNAs exceeding the number of ribosomes 10-fold (Waldron and Lacroute, 1975). The high numerical abundance of tRNA molecules strengthens our hypothesis that the doubling of 275 highly-transcribed tRNA genes results in a cellular burden leading to a compensatory ploidy increase. Furthermore, recent work suggests that increases to ploidy in *S. cerevisiae* is a relatively common phenomenon, especially under conditions of stress (Harari *et al*., 2018a; Harari *et al*., 2018b). We postulate that haploid cells are less amenable to compensate to this tDNA overabundance, with this conclusion supported by an apparent improvement in tRNA neochromosome stability in diploid cells. Despite our inconclusive results when attempting to downregulate tRNA expression through a constitutively active *MAF1,* our observations following transcriptomic and proteomic characterization further lend credence to the hypothesis that a relative doubling in the number of tRNA genes in haploids leads to an overload of uncharged tRNAs and thus a dysregulation of the translational machinery. A higher ratio of uncharged:charged tRNAs may interfere with protein biosynthesis and potentially cause a general stress response.

Secondary observed phenomena include structural variations of the tRNA neochromosome and its unstable nature. Given the highly-transcribed nature of tRNA genes, a general instability of the tRNA neochromosome is not unexpected. We postulate that the complex structural variations may be related to its circular intermediary structure, due to the known instability of ring chromosomes in higher eukaryotes (McClintock, 1938), and potential for complex variations (including duplications, deletions or inversions) caused by dicentric or catenane formation (Guilherme *et al*., 2011; Pristyazhnyuk and Menzorov, 2018). Furthermore, its logical to assume that a general selection pressure exists in haploid cells against an intact tRNA neochromosome structure, ultimately leading to a reduction in its overall size.

Although the tRNA neochromosome conferred an expected cellular burden, we have not been able to pinpoint a singular cause, suggesting that its origin may be multifactorial. Aside from an overabundance of tRNA genes, other potential sources include perturbation of tRNA neochromosome segregation during cell division leading to daughter cell aneuploidies. Ultimately, elucidation and characterization attempts will greatly benefit once the tRNA neochromosome is introduced into the final synthetic yeast genome as part of Sc2.0, absent of its native tRNA copies.

tRNA sequencing analysis demonstrate that the bulk of the tRNA genes of the tRNA neochromosome are expressed in a manner comparable to their wild-type counterparts. However, a balancing effect on the cellular tRNA pool can be observed, suggesting an overall regulation or buffering effect within the cell, or alternatively a competition for RNAPIII. Notably, earlier examples have explored mechanisms with the specific tRNA pool alteration to specific stress responses to match the codon usage of genes required for the cellular response (Torrent *et al*., 2018). Although we previously identified a potential tRNA_Ser_^CGA^ maturation deficiency (Zhao *et al*., 2022), our global tRNAseq analysis could not identify evidence of any global processing defects. It is possible that the structure of the 5’ and 3’ leader and trailer sequences influence the processing efficiency of RNase P: previous work has shown that complementary pairing between the 5’ and 3’ sequences affects RNase P cleavage, and that these sequences may need to be separated for substrate recognition (Ziehler et al., 2000).

As a designer biological structure, the tRNA neochromosome provides a unique insight into how such entities function within the cell. Replication profiling provided evidence of replication fork firing with associated troughs located near replication termination (*TER*) sites. Interestingly, nucleosome mapping data demonstrates that tRNA expression may contribute to the positioning of the nucleosomes. Nucleosome positioning differs slightly compared to that found in the genome, with a shorter linker between individual nucleosomes presumably caused by a higher overall density and shorter distances between tDNAs than are observed in the native genome. Furthermore, FISH and Hi-C provides conclusive evidence that the tRNA neochromosome does not co-localize with the nucleolus, supporting the tethering hypothesis proposed recently (Belagal *et al*., 2016). Finally, we expect that the Dre-*rox* recombination system will be a powerful future system to address questions of tRNA genetics in a systematic manner, especially once the fully synthetic Sc2.0 strain is available.

We recognize that, although our design attempted to balance and reconcile risk, function and stability, our approach was inherently arbitrary based on then-current knowledge of chromosome and tRNA biology. Despite these limitations, improved knowledge of chromosome biology will enable the design and construction of new and improved tRNA neochromosome variants in future to meet the objectives of the Sc2.0 project. This principle is important as part of any Design-Build-Test-Learn cycle and may be important to maximize fitness of the final synthetic strain. For example, more recent evidence suggests that some tRNA introns may indeed have a physiological role (Hayashi *et al*., 2019), and so future variants will likely benefit from their presence.

Overall, this study has shown the remarkable tolerance of baker’s yeast to support the presence of an additional neochromosome housing 275 tRNA genes. Looking beyond the Sc2.0 project, this work demonstrates the application of these designer structures and the benefits of radically re-engineering the host cell machinery. Future designer neochromosomes will provide a means to address fundamental questions of biology through the refactoring of genetic components in a systematic manner (Postma *et al*., 2021), introduce novel characteristics for designer industrial strains (Kutyna *et al*., 2022), provide a chassis to incorporate large-scale metabolic pathways and provide the means to further understand aspects of chromosome biology through a ‘build to understand’ approach. Given rapid technological advances and reduced costs of DNA synthesis, we anticipate the routine construction of custom neochromosomes leading to tangible applications in the near future.

## Data and code availability

- WGS data, tRNAseq data, RNAseq data, nucleosome positioning data and Hi-C data is deposited at SRA with BioProject number PRJNA884518 and are publicly available as of the date of publication.
- Proteome data are deposited at PRIDE and will be publicly available as of the date of publication.
- Code for assigning tRNA genes to flanking sequences is deposited at GitHub and will be publicly available as of the date of publication.
- Any additional information required to reanalyze the data reported in this study is available from the lead contact upon request.

## Acknowledgments

This work was supported by UK Biotechnology and Biological Sciences Research Council (BBSRC) grants BB/M005690/1, BB/P02114X/1 and BB/W014483/1, and a Volkswagen Foundation “Life? Initiative” Grant (Ref. 94 771) to YC. DS is supported by the Max Planck Society in the framework of the MaxGENESYS project. RSKW, at the University of Edinburgh, was supported by EPSRC studentship (1419736) and the Bill and Melinda Gates Foundation. RSKW, as part the *Yeast 2.0* initiative at Macquarie University, is financially supported by Bioplatforms Australia, the New South Wales (NSW) Chief Scientist and Engineer, and the Australian Research Council (ARC) *Centre of Excellence in Synthetic Biology*. FA is supported by a grant from the Spanish Ministry of Science and Innovation (PID2020-118423GB-I00). CAN acknowledges support from the Biotechnology and Biological Sciences Research Council (BBSRC), part of UK Research and Innovation, Grant BB/N016858/1, Core Capability Grant BB/CCG1720/1 and the National Capability (BBS/E/T/000PR9814). JD is supported by a Royal Society Newton Advanced Fellowship (NAF\R2\180590) hosted by YC. JDB was supported by NSF grants MCB-1026068, MCB-1443299, MCB-1616111 and MCB-1921641.

## Author Contributions

Conceptualization and coordination (JDB, YC). Designed and constructed the tRNA neochromosome (DS, RSKW, YL, JDB, YC). Designed experiments (DS, RSKW, FA, YS, CAN, RK, JD, LMS, JDB, YC). Conducted experiments (DS, RSKW, SJ, ANB, YW, CAM, CC, AG, DS2, JM, LO, WL, KJ, DA, EGR, TWT, RS). Data analysis (DS, RSKW, ANB, YW, CAM, CC, AG, DS2, JM, JC, BAB, TWT, TE, FA, YS, CAN, RK, JD, LMS, JDB, YC). Manuscript writing (DS, RSKW, YC). All authors revised and approved the final manuscript.

## Material and Methods

### Yeast growth media and cultivation

All yeast strains were derivatives of *S288C* and grown at 30°C unless otherwise specified. A full list of strains used may be found in Table S1. Yeast cells were grown in either YEP/YPD media (10 g/L yeast extract, 20 g/L peptone and with or without 0.64 g/L tryptophan) or Synthetic Complete (SC) media lacking the indicated amino acids, both supplemented with 2% glucose if not stated otherwise. Standard solid media contained 2% agar and cultures were incubated at 30°C. Liquid cultures were cultivated according to standard procedures in glass tubes on a TC-7 roller drum (New Brunswick).

### Yeast transformation

Yeast transformation was undertaken according to a modified protocol described by Daniel Gietz and Woods (2002). Briefly, yeast strains were inoculated into 10 mL of liquid media and incubated with rotation at 30°C overnight. Overnight cultures were then re-inoculated to OD_600_ = 0.1 and incubated with rotation to a target OD_600_ of between 0.5 and 1. Cells were then centrifuged at 2,103 g for 5 minutes, washed with 10 mL sterile ddH_2_O and centrifuged again. Cell pellets were washed with 10 mL of 0.1 M LiOAc, centrifuged at 2,103 g for 5 minutes. 50 μL of cells were mixed with 5 to 10 μL of transforming DNA, 36 μL of 1M LiOAc, 19 μL ddH2O, 25 μL 10 mg/mL herring sperm carrier DNA and 240 μL of 44% PEG 3350. Transformations were incubated at 30°C for 30 minutes before adding 36 μL of DMSO and heat-shock at 42°C for 15 minutes. Cells were pelleted and incubated in 400 μL of 5 mM CaCl_2_ at room temperature for 10 minutes and plated onto selective media.

### Automating flanking sequence assignment to tRNA genes

An algorithm (Fig. S1A) and associated scripts (https://github.com/cailab-uom/neochromosome) were used to assign tRNA genes to respective flanking sequences recovered from the *A. gossypii* and *E. cymbalariae* genome. These were based on custom Python scripts with BioPython modules. *S. cerevisiae* tRNA gene sequences (lacking introns), and *A. gossypii*/*E. cymbalariae* 5’ and 3’ flanking sequences were recovered from Genbank files downloaded from the NCBI genome database (accession numbers may be found in Tab. S8), with metadata exported to comma separated value (.csv) files. All tRNA gene anticodons were determined (*i.e.* independently verified) by ‘calling’ a program compiled from the source code of tRNAscan-SE (Schattner *et al*., 2005). We developed an algorithm to preferentially map tRNA genes to flanking sequence based on the following design parameters: (1) tRNA genes of *S. cerevisiae* should match flanking sequences in *A. gossypii* and *E. cymbalariae*, in that order of preference, based on relative orientation and matching anticodon where possible. (2) Flanking sequences should only be assigned to tRNA genes once. (3) tRNA genes will lack introns. (4) All 3’ tRNA flanking sequences should possess five or more thymidine residues required for transcriptional termination. (5) The 500 bp 5’ flanking sequences should not contain unwanted features. (6) *rox* recombination sites will flank each tRNA cassette. To address the 5^th^ design parameter above, scripts were used to remove and alter unwanted features in tRNA flanking sequences (summarized in Fig. S1B), and include the translational silencing of gene starts, avoidance of those with overlapping regions and the removal of Solo LTR sequences for *E. cymbalariae*. Some tRNA genes were discarded from the available pool altogether. Following tRNA gene/flanking sequence assignment, tRNA cassettes were concatenated into tRNA arrays and generated as Genbank output files. Table S3 details a full list of each *S. cerevisiae* tRNA gene and its associated assigned flanking sequence. Supplementary Data S1 includes the output file generated from scripts, including annotated tRNA arrays and metadata on tRNA genes and their associated flanking sequences. tRNA gene flanking sequences were also obtained from the yeast, *Kluyveromyces lactis*, but were not used in our final design.

### Manually curated design differences

The *chrVI* tRNA array was manually designed prior to the development of custom scripts. In this instance, 20 bp 3’ flanking sequences from *A. gossypii* were incorporated (with the exception of the tG(GCC)F1 (SC.t6.4) tRNA cassette, which utilized a 40 bp 3’ flanking sequence to capture the full poly-T sequence). Furthermore, the *chrVI* tRNA retains SfiI restriction sites which were used to experimentally ‘flip’ two tRNA cassettes from their wild-type orientation and ensure unidirectionality. For this reason, the orders of the tG(GCC)F1 (SC.t6.4) and tY(GUA)F1 (SC.t6.5) tRNA cassettes are reversed. Finally, following manual sequence validation of the tRNA neochromosome following flanking sequence assignment, a further four tRNA cassettes were observed to house unwanted features and were thus manually assigned different flanking sequences. These tRNA genes included *tV(AAC)G1* (SC.t07.12), *tV(AAC)H* (SC.t08.02), *tL(UAA)J* (SC.t10.17) and *tL(UAG)L2* (SC.t12.12) (Tab. S3).

### Neochromosome design

The tRNA neochromosome itself was manually designed *in silico* from constituent tRNA arrays using Snapgene (Chicago, USA) and was used as a reference point for the design of eSwAP-In vectors used in its construction. The designed and sequenced tRNA neochromosome sequences are deposited at NCBI BioProject PRJNA351844.

### Origins of replication and design of replication termination sites

ARS elements were obtained from *Candida glabrata* (*chrF-444*, *chrL-615* and *chrM-794*; sequences listed in Tab. S7). These were chosen on the basis of an intergenic ORC-binding motif that closely resembles the motif found in all *S. cerevisiae* origins. The ARS *chrM-794* origin of replication was later observed to unexpectedly house an unannotated tRNA gene from *Candida glabrata* (revealed to be *chrM*.trna11-GlyGCC). Bidirectional replication termination sites (sequences listed in Tab. S7) were designed to contain two conserved Fob1 recognition sites as follows: ***TERI***: A hybrid between the *TER2* and *TER1* sites of *Saccharomyces paradoxus* and the *TER1* and *TER2* sites of *Saccharomyces uvarum*. ***TERII*:** A hybrid between the *TER2* and *TER1* sites of *Saccharomyces kudriavzevii* and the *TER1* and *TER2* sites of *Saccharomyces pastorianus*. ***TERIII*:** A hybrid between the *TER2* and *TER1* of *Saccharomyces mikatae* and the *TER1* of *Saccharomyces paradoxus* with a modified *TER2* hybrid containing a flipped middle spacer region. ***TERIV*:** A hybrid between the *TER2* of *Saccharomyces paradoxus* and the *TER1* site of *Saccharomyces uvarum* as well as the *TER1* of *Saccharomyces kudriavzevii* and *TER2* of *Saccharomyces pastorianus* (with some minor modifications). *TER* sites *I-IV* were synthesized by Twist Biosciences (San Francisco, USA).

### Universal homologous region

The Universal Homologous Region (UHR) was designed using a Perl script to produce a sequence of randomly-generated DNA 500 bp in length. This sequence was subjected to BLAST (https://blast.ncbi.nlm.nih.gov/Blast.cgi) against the *S. cerevisiae* genome, with no significant homology found. This DNA sequence is listed in Table S8. The UHR was synthesized *de novo* from overlapping oligonucleotides using the overlap-extension PCR method from the Build-A-Genome course (Cooper *et al*., 2012).

### Assembly of tRNA arrays

The *chrI* tRNA array was synthesized *de novo* from overlapping oligonucleotides into four ‘building blocks’ containing 100 bp overlapping regions using the overlap-extension PCR method from the Build-A-Genome course (Cooper *et al*., 2012) and assembled using Gibson Assembly before verification with Sanger sequencing (Edinburgh Genomics, Edinburgh, UK). tRNA arrays over 10 kb in size were initially synthesized as sequences less than 10 kb and later combined into a linearized pRS413 acceptor plasmid using either Gibson Assembly (Gibson *et al*., 2009) or *in vivo* homologous recombination in BY4741 using the LioAC transformation method (Gietz and Woods, 2002). To facilitate assembly, 100 bp overlapping region was incorporated into the central region of each split tRNA array. tRNA arrays (listed in Tab. S2) were synthesized *de novo* by Wuxi Qinglan Biotech Co. Ltd (Yixing City, China), except for the *chrVIII*, *chrIX* and *chrXVI* tRNA arrays that were synthesized by Thermo Fisher Scientific (Renfrew, UK). Once constructed, tRNA genes were excised from their respective backbone using terminal restriction sites. All DNA constructs were verified using standard methods (colony PCR, restriction digest fingerprinting and Sanger sequencing).

### Neochromosome construction (eSwAP-In)

We developed an approach (later referred to as eSwAP-In) to construct the tRNA neochromosome, although this approach differs in key aspects to that described by Mitchell *et al*. (2021) and are noted as follows. tRNA arrays were introduced into specialized eSwAP-In vectors to facilitate construction. These vectors housed genetic elements required for the eSwAP-In process, including both the *LEU2* or *URA3* auxotrophic markers and a 500 bp ‘Universal Homologous Region’ (UHR). tRNA arrays were excised from their backbone using flanking restriction sites (Tab. S5), with eSwAP-In vectors linearized using the SmaI restriction enzyme. tRNA arrays were inserted into their respective eSwAP-In vector using Gibson Assembly or *in vivo* homologous recombination in BY4741 facilitated by two homologous 330 bp to 650 bp “bridge” sequences amplified using PCR (*cf*. Fig. S2A). These bridge sequences also introduced functional elements to the growing neochromosome, including the origins of replication and the bidirectional *TER* sites (*cf*. Fig. S2B). The primers used to amplify each bridge sequences were designed to house the necessary 40 bp overhangs for subsequent assembly. Neochromosome construction was performed in a stepwise manner (*cf*. Fig. 2A) using repeated rounds of *LEU2* and *URA3* marker swapping in BY4741 and *synIII/VI/IXR*. Each tRNA array and sequences necessary for eSwAP-In were first recovered from their backbone using flanking NotI or FseI restriction enzymes and transformed into yeast containing the growing circular neochromosome. Candidate isolates were re-streaked onto selective agar for single colonies to induce loss of any prior auxotrophic marker before replica-plating onto agar lacking histidine (SC-His), leucine (SC-Leu) and uracil (SC-Ura), respectively, to determine the correct phenotype. Verification of the intact neochromosome was then performed using the PCR Tags (Tab. S4). Finally, to facilitate the integration of the telomerator cassette for subsequent linearization, the residual *URA3* marker was ‘popped-out’ on 5-FOA following the final round of tRNA neochromosome construction. This was undertaken by introducing a second homologous 500 bp UHR located upstream of the *URA3* marker (introduced as a PCR ‘bridge’ during construction of the eSwAP-In vector) (*cf*. Fig. S2B).

### PCR Tags method and conditions

To screen for candidate isolates, primers were designed to anneal to the 5’ and 3’ flanking sequence of each tRNA cassette to produce an amplicon of approximately 400 bp to 600 bp. After each round of eSwAP-In, an initial set of one to six primer pairs were used to screen colonies for an intact tRNA neochromosome before undertaking a full set of PCR Tags for all 275 tRNA cassettes. Crude genomic DNA was generated by incubating yeast isolates at 95°C in 50 μL of 20 mM NaOH for 10 minutes or the MasterPure™ Yeast DNA Purification Kit (Lucigen). PCR Tag analysis was performed either by adding 1.8 μL of yeast lysate used as template DNA to 6.25 μL of 2X GoTaq® Green Master Mix (Promega), 4.75 μL of nuclease-free water and 400 nM of each primer or by adding 1 µl of yeast lysate to 5 µl of DreamTaq Green PCR Master Mix (2X) (ThermoFisher Scientific) plus 4 µL of nuclease-free water with 200 nM of each primer. PCR reactions were performed as follows: 95°C for 2 min followed by 30 cycles of 95°C /2 min 30 s, 50°C /1 min 15 s, 72°C /1 min 45 s followed by a 72°C /5 min extension time (GoTaq) or 95°C for 2 min followed by 40 cycles of 95°C /20 s, 52°C /45 s, 72°C /1 min 30 s followed by a 72°C /2 min extension time (DreamTaq). 8-10 μL aliquots of each reaction were then run on a 1.5-2% TAE agarose gel. A full list of PCR Tag primers used may be found in Table S4; a representative example gel may be found in Figure S2D.

### Repair of tRNA neochromosome structural variations

The structural variation observed in YCy1576 was repaired as follows. Initially, *URA3* and *LEU2* markers, each coupled with an I-*Sce*I recognition site, where simultaneously integrated upstream and downstream of the structural re-arrangement, deleting the duplicated part of the *chrXIII* tRNA array and the partially complete *chrX* tRNA array (YCy2411). In the second round of repair, two plasmids were co-transformed. The first plasmid (*KANMX* marker) contained the *chrX* tRNA array flanked by I-*Sce*I recognition sites; the second (*MET15* marker) housed a galactose inducible I-*Sce*I overexpression construct. Subsequent induction of I-*Sce*I expression facilitated the release of the *chrX* tRNA array from the first plasmid and generation of two neochromosome fragments. The presence of 500 bp homologous sequences on the three fragments facilitated recombination-mediated integration of the *chrX* tRNA array and the repair of the inversion to generate the final tRNA neochromosome (YCy2508) (*cf*. Fig. 2B). Single colonies were selected on SC-His-Met + 5-FOA with 2% galactose agar plates. Candidates were screened with primers to detect the correct homologous recombination borders as well as the missing tRNA cassettes of the *chrX* tRNA array before full PCR Tag analysis, size verification *via* PFGE and short read sequencing was performed resulting in the final strain YCy2508. The raw reads of YCy2508 are deposited at NCBI BioProject PRJNA351844 (SAMN30265401).

### Whole genome sequencing

Yeast genomic DNA was prepared from 5 mL stationary phase culture either with the MasterPure™ Yeast DNA Purification Kit (Lucigen) according to the manufacturer guidelines or using the Cetyl Trimethyl Ammonium Bromide (CTAB) DNA extraction method. Briefly, cells were harvested at 4,000 g for 4 min, resuspended in 1 mL extraction buffer (2% CTAB, 100 mM Tris, 1.4 M NaCl and 10 mM EDTA, pH 8.0) and transferred into a 2 mL screw-top tube containing ∼50 mg 425–600 μm washed glass beads (Sigma). Cells were lysed by vortexing for 10 min at maximum speed followed by incubation for 10 min at 65°C. Cell debris were removed by centrifugation at maximum speed for 2 min and supernatant was transferred to a 2 mL tube, 4 μL of 100 mg/mL RNase (Qiagen) was added, followed by incubated at 37°C for 15 min. 700 μL of chloroform was added, mixed carefully and centrifuged at maximum speed for 2 min. The aqueous phase was transferred into a 1.5 mL reaction tube containing ∼0.6 volumes of isopropanol. Thorough mixing was performed with the precipitate collected by centrifugation at maximum speed for 10 min. The resulting supernatant was discarded and the pellet was washed with 500 μL 70% ethanol followed by centrifuging for 5 min at maximum speed. The supernatant was decanted and the tube was again pulse-centrifuged with the residual alcohol removed with a pipette followed by air drying for 2-5 min. The pellet was subsequently resuspended in 100 μL of H_2_O. DNA library preparation was performed using the Nextera XT kit (Illumina). Tagmentation, inactivation, and library amplification were performed following kit instructions and using commercial reagents. Amplified libraries were purified using 1.8× volumes AMPure XP beads and eluted in 10 µl nuclease-free water. Library yield was quantified using the Qubit high-sensitivity DNA kit and library quality was determined on an Agilent Bioanalyzer (high-sensitivity DNA assay). Samples were pooled and 150 bp paired-end reads were generated on an Illumina NextSeq 500. Alternatively, PCR-free whole genome sequencing was performed using the Sanger Institute service. Samples were normalized so that 500 to 1000 ng of input DNA enters the process. DNA was fragmented to 450 bp using a Covaris LE220 sonicator prior to library construction. During library construction fragmented DNA was purified ready for End-repair, A-tailing and unique dual indexed adapter ligation with no amplification step. Libraries were quantified using qPCR and then pooled and 150 bp paired-end reads were generated on an Illumina HiSeqX. Sequencing reads QC, data processing and analysis were performed using standard procedures, clean reads were mapped to the respective reference sequence using Bowtie2 v2.3.5.1 with default parameters (Langmead *et al*., 2009). GATK v2.7 (McKenna *et al*., 2010) and SAMtools v0.1.19 (Li *et al*., 2009) were used in conjunction with IGV for analysis.

### Chemical extraction and transfer of the tRNA neochromosome

Extraction and transfer of circular neochromosome variants from one cell to another was undertaken according to the method described by (Noskov *et al*., 2011). The Plasmid Maxi Kit (Qiagen) with QS solution (1.5 M NaCl, 100 mM MOPS, 15% v/v isopropanol) and PlasmidSafe (Epicentre) were used *in lieu* of the Qiagen Large Construct Kit and Qiagen exonuclease, respectively. Purified neochromosome DNA was then transformed into BY4741 and BY4742 using the spheroplast transformation method described by Burgers and Percival (1987). Increased volumes were used during transformation, with 12 μL of neochromosome DNA added to 300 μL of spheroplasts. Transformed spheroplasts were then added to 10 mL of molten SC-Ura or SC-His TOP agar (SC-Ura or SC-His media containing 1 M sorbitol and 2.5% agar; held at 45°C), gently mixed, and poured directly onto petri dishes containing 10 mL of SC-Ura or SC-His SORB (SC-Ura or SC-His media containing 0.9 M sorbitol, 2% agar and 3% glucose). The plates were then incubated at 30°C for four days before the appearance of sufficiently large colonies.

### Neochromosome linearization

Telomere seed sequence integration and neochromosome linearization was performed according to the method described by Mitchell and Boeke (2014). Following the integration of the telomerator cassette into defined regions of the neochromosome, strains were transformed with a plasmid expressing the I-*Sce*I homing endonuclease under a control of a *GAL1* promoter. Isolates were then inoculated into 10 mL of SC-His-Ura-Leu (2% glucose) and incubated with rotation at 30°C overnight. Once saturated, cultures were washed once with sterile PBS (phosphate buffered saline) solution and re-inoculated into 10 mL of SC-His-Leu containing either 2% galactose (for induction of I-*Sce*I) or 2% glucose (negative control) to a target OD_600_ of 0.05 and incubated with rotation at 30°C for 24 hours. After 24 hours of growth, cultures were serially-diluted by a factor of 1 in 1,000 and plated onto SC-His + 5-FOA (1 mg/mL). Approximate efficiency of linearization was inferred by comparing the number of colonies of galactose-induced cultures with those cultured in glucose (normalized for cell density). After three days of growth, yeast isolates were verified using a colony PCR with primers designed to anneal with each arm of the telomerator cassette and to screen for the presence or absence of the junction region at the I-*Sce*I cut site.

### Pulsed field gel electrophoresis (PFGE)

Plug preparation and pulsed-field gel electrophoresis (PFGE) was performed according to previously described methods by Hage and Houseley (2013). PFGE was undertaken by running samples on a 1.3% agarose gel in 0.5 X TBE solution at 14°C on a Bio-Rad clamped homogenous electric field (CHEF) apparatus (CHEF-DR III). 6 V/cm were used with a ramped switch time of 15-25 s for 24 h. Alternatively, samples were run for 15 h with a switch time of 60 s, followed by a further 15 h with a switch time of 120 s at a 120° angle. The resulting gel was stained with 1 X SYBR Safe (ThermoFisher Scientific) and imaged using a Fujifilm FLA-5100 or Typhoon Trio (GE) laser scanning system. The identity and size of the tRNA neochromosome was inferred by lambda ladders run on the same gel and/or the known molecular karyotype of BY4741.

### tRNA neochromosome stability assay

Overnight cultures of haploid BY4741, BY4742 and diploid *MAT***a**/**a** and *MAT***a**/α containing the tRNA neochromosome (YCy2837, YCy2838, YCy2508 and YCy2844) were inoculated as biological triplicates into 20 mL SC-His to a starting OD_600_ of 0.1. Every 24 h, cultures were back diluted to OD_600_ 0.1 and glycerol stocks were prepared over a time-course of seven days corresponding to approximately 50 generations. On the seventh day, dilution series were plated on SC-His media and four randomly selected candidates per strain were tested with a subset of the PCR Tag primers via colony PCR. DNA of 2 mL overnight culture was extracted with the MasterPure™ Yeast DNA Purification Kit (Lucigen) and whole PCR Tag analysis was performed on population level to test the general presence of all tRNAs within the population (data not shown). Selected candidates were submitted to Nanopore sequencing to identify the location of recombination.

### Nanopore sequencing

High-molecular weight DNA extraction was performed according to a modified Qiagen Genomic-tip 100/g protocol. Yeast cultures were grown to early stationary phase (OD_600_ of approximately 10) and an equivalent OD_600_ of 250 were harvested with centrifugation at 5000 RCF. Spheroplasts were generated by incubating resuspended yeast pellets in 4 mL of Y1 buffer supplemented with 250 U lyticase (Sigma Aldrich) at 37°C for 1 hour. Spheroplasts were then harvested at 800 RCF for 10 minutes at 4°C. The pellet was resuspended in G2 Buffer with 20 µL RNase A (100 mg/ml) (Qiagen) and 125 µL RNase Cocktail (Invitrogen). After incubation for 5 min at RT, 100 µL Proteinase K (Sigma Aldrich) was added and incubated for 1 hour at 50°C. The lysate was cleared by centrifuging at 5000 RCF for 15 min at 4°C before loading onto the column. The remaining procedure was performed according to the manufacturer’s guidelines. DNA quality was accessed via gel-electrophoresis, NanoDrop™ 2000 spectrophotometer and Qubit 4 fluorometer using the dsDNA BR reagents. Library preparation was performed using the SQK-LSK109 and the Native Barcoding kits (EXP-NDB104 and EXP-NDB114) according to the manufacturing guidelines. The only alteration was to increase the DNA mass 5-fold to match the required starting material molarity, DNA is approx. 50 kb in size based on our initial PFGE analysis (data not shown). Sequencing was performed on a MinION Mk1B device (Oxford Nanopore Technologies) using flow cell chemistry R9.4.1 (FLO-MIN106D). Base calling was performed using Guppy and minimap2 was used for sequence alignments. Samtools and IGV were used for data analysis.

### Tecan growth curve assay

Growth curve analysis was performed in a Tecan F200 plate reader in 96-well flat bottom plates (Corning) with 150 µL of medium inoculated to a starting OD_600_ of 0.05. Plates were sealed with TopSeal-A PLUS (Perkin Elmer). The following kinetic parameters were run with a varying number of cycles (289 cycles for approx. 24 hrs at 30°C): 2:23 min orbital shaking (1.5 mm amplitude), 2:23 min linear shaking (2 mm amplitude) followed by OD measurement at 595 nm. Data was analyzed and visualized with custom R scripts and the R package Growthcurver (Sprouffske and Wagner, 2016).

### Single-copy, essential, tRNA gene complementation

To assay synthetic tRNA functionality on an individual level, three, single-copy tRNA genes were investigated: *SUP61 (tS(CGA)C)*, *TRT2 (tT(CGU)K)* and *TRR4 (tR(CCG)L)*. All three tRNA genes are essential in *S. cerevisiae*. Synthetic variants of each were integrated into the *HO* locus of BY4741, with *URA3* used as a selective marker for integration. Wild-type copies of each essential tRNA gene were removed by PCR-amplifying a *KanMX* marker with 40 bp homology arms specific to the surrounding region of each tRNA gene (viability was supported by episomal copies of each). Removal of wild-type tRNA genes and integration into the *HO* locus was verified using colony PCR, with primer pairs designed to amplify the junction regions of each locus. Following induced loss of each respective episomal copy, biological duplicates of these strains (YCy896 to YCy905) and a wild-type control (BY4741 + pRS416) were inoculated into 10 mL of SC-Ura and incubated with rotation overnight at 30°C. Overnight cultures were then adjusted to a normalized OD_600_ of 0.03 in sterile PBS (phosphate buffered saline) solution, serially diluted, and spotted side-by-side onto SC-Ura selective media (Fig. S1C). Plates were then incubated at 30°C for three days.

### Dre recombinase-induced tRNA copy number adjustment: proof-of-concept

A proof-of-concept with Dre recombinase was performed on the *chrVI* tRNA array in BY4741. The *chrVI* tRNA array contained 10 tRNA genes from *chrVI* and one tRNA gene from *chrV*. The pRS413-*chrVI*-tRNA array was first transformed into BY4741, followed by a second transformation with pRS415-P_SCW11_*DreEBD* (Liu *et al*., 2018b). The double transformed strain was cultured in SC-His-Leu liquid medium overnight and re-inoculated to OD_600_ of 0.1 into two tubes, one with a 7 h induction of 1 μM β-estradiol and the other un-induced. Following induction, cells were plated onto SC-His, with single colonies picked for plasmid extraction and recovery in *E. coli*. XbaI restriction sites were subsequently used to excise the *chrVI* tRNA array from its plasmid backbone and generate a restriction map indicating the relative reduction in the number of tRNA cassettes. Sanger sequencing was then used to determine the full genotype information of the remaining tRNA genes.

### Ploidy determination by flow cytometry

The ploidy of yeast cells was determined using SYTOX Green stained fixed cells in a SH800S Cell Sorter (Sony Biotechnology). Briefly, cells were grown to an OD_600_ of approx. 0.5 with 2 mL of cells collected (4 min at 2000 RCF), washed with filtered H_2_O and fixed in 1 mL 70% EtOH. Fixed cells were washed twice in 1 mL filter sterilized 50 mM sodium citrate followed by resuspension in 1 mL 50 mM sodium citrate with RNase A (final conc. 0.25 mg/mL) and incubation at 50°C for 1 hr. Subsequently Proteinase K was added with a final conc. of 0.4 mg/mL followed by 1 hr at 50°C before harvesting the cells (4 min 2000 RCF). Cell pellets were resuspended in 1 mL 50 mM sodium citrate with Sytox Green (5 mM stock; 1:5000 diluted) and subsequently measured in the SH800S Cell Sorter. Known haploid (BY4741) and diploid (BY4743) control strains were always used as standards.

### Fluorescent in situ hybridization (FISH)

Yeast strains (YCy2508, YCy2649) were cultured in 3 mL of SC-His overnight at 30°C before back-diluting into 5 mL fresh SC-His (OD_600_ = 0.1), and subsequently grown to a concentration of approx. 2*10^7^ cells per mL. Slides and hybridization of probes were prepared according to Scherthan and Loidl (2010). Hybridization reactions were performed using 6 μL of a 3 μM 5’-Cy3 labelled probe (5’-taactttaaataatgccaattatttaaagtta-3’) for the rox recombination sites, and 3 μL the three 3 μM 5’-FAM labelled probes, targeting the *5S* rDNA according to Thompson *et al*. (2003), for rDNA sequences, respectively. Slides were imaged using a Nikon A1 confocal microscope with a 60x objective. Three different derivatives of YCy2508 which lost different PCR Tags were used for the imaging with n = >100 cells for each derivate. Results were merged and used to calculate the percentage of cells with different localization patterns.

### Live cell imaging

Yeast strains (JSY396 and JSY397) were cultured into 3 mL of SC-His overnight at 30°C before back-dilution into 5 mL fresh SC-His (OD_600_ = 0.15) and grown 5 hours prior imaging. Cells were harvested and washed with H_2_O. Live cell imaging was performed on a Nikon A1 confocal microscope with a 60x objective. The recognition sequence of the constructed TAL-GFP is 5’-ttaaataattggcattat-3’. The results of four clones of JSY396 were used to calculate the percentage of cells with different patterns. More than 250 cells for each clone were counted.

### RNA-seq analysis

The strains with 3 biological replicates were prepared using sample preparation methods established previously for transcriptome analysis (Shen *et al*., 2017a). The total RNA was prepared using the Omega Bio-tek Yeast RNA Kit (CAT# R6870-02). A 200-400bp RNA-seq library was prepared by the MGIEasy^TM^ RNA Library Prep Kit V3.0 (CAT# 1000006384). The transcriptome sequencing was performed with the BGISEQ-500 platform. For transcriptomic analysis, reads were mapped to genome sequence by hisat2 v2.1.0 (Kim *et al*., 2019), assigned read counts for each gene by featureCounts v2.0.1 (Liao *et al*., 2014) and differentially expressed genes were analyzed by DEseq2 v1.30.1 (Anders and Huber, 2010). Genes were assessed for significantly differential expression if the log_2_(Foldchange) was >1 or <-1, and if the raw P value fell below the threshold of the 5% Family Wise Error Rate (FWER) after Bonferroni correction (threshold = 7.62E-6). To identify differentially active biological processes between tRNA neochromosome strains and its native control strains, gene enrichment analyses were performed as described previously (Shen *et al*., 2017a) using yeast KEGG pathways and yeast Gene Ontology (GO) annotations. The significance of each KEGG pathway and GO term in genes was individually identified using the hyper-geometric test and Chi-squared test the threshold P-value <0.001.

### Proteome analysis

The strains with 3 biological replicates were prepared using sample preparation methods established previously for proteome analysis (Shen *et al*., 2017a). Yeast protein was extracted with Urea, reduced, alkylated, and digested with trypsin (Shi *et al*., 2021). The protein identification and quantitation performed by tandem mass spectrometry. The proteins were labeled by Thermo Scientific TMT pro 16plex Label Reagent Set (CAT# A44522) according to the manufacturer’s instructions, and then the labeled peptides were fractionated with high pH reversed-phase (RP) chromatography method (Wang *et al*., 2011) and analyzed by a Q Exactive^TM^ HF-X mass spectrometer (Thermo Fisher Scientific, San Jose, CA) coupled with an online High-Performance Liquid Chromatography (HPLC, Thermo Scientific™ UltiMate™ 3000 UHPLC system). For proteomic analysis, Mascot and IQuant (Wen *et al*., 2014) were used for protein identification and quantification. The differential expression proteins were identified if the log_2_(Foldchange) was >1 or <-1, and the P value <0.001. The Gene enrichment analyses of KEGG pathways and Gene Ontology were performed as above description.

### Preparation of mononucleosomal DNA and generation of nucleosomal occupancy maps

*S. cerevisiae* cultures of 200 mL at 0.8 x 10^7^ cells/mL were collected for preparation of mononucleosomal DNA. Cells were treated with 10 mg of Zymolyase 20T for 5 minutes at 30°C to generate spheroplasts. Mononucleosomal fragments were generated by digesting DNA with 300 units/mL of micrococcal nuclease (MNase) at 37°C for 10 minutes. The amount of MNase was optimized experimentally to generate an 80:20 ratio of mononucleosomes to dinucleosomes, as described by González *et al*. (2016); Soriano *et al*. (2014). Mononucleosomal DNA was recovered from 1.5% agarose gels to prepare sequencing libraries following the Illumina protocol. They were sequenced in an Illumina NextSeq 500 platform using the paired-end protocol. Reads were aligned using Bowtie (Langmead *et al*., 2009) to the *S. cereviase* (SacCer 3) genome or to the tRNA neochromosome sequence. Alignment files were processed using the NUCwave algorithm (Quintales *et al*., 2015) to generate the nucleosome occupancy maps.

### Hi-C analysis

Hi-C experiments were performed following the protocol described in Lazar-Stefanita *et al*. (2017) and adapted in Dauban *et al*. (2020). Briefly, 1 to 3 x 10^9^ cells in 150 mL synthetic medium were crosslinked using 3% formaldehyde. Quenching of formaldehyde was performed using 300 mM glycine at 4°C for 20 min. Fixed cells were recovered through centrifugation, washed, and stored at −80°C. The Hi-C library was generated using a DpnII restriction enzyme as described in Lazar-Stefanita *et al*. (2017), except that after the protein digestion step the DNA was purified using AMPure XP beads. The DNA Hi-C libraries were sheared into 300 bp using a Covaris S220 apparatus (Covaris) and the biotin-labeled fragments were selectively captured by Dynabeads Myone Streptavidin C1. The sequencing library was generated using Invitrogen Collibri PS DNA Library Prep Kit for Illumina, following manufacturer instructions.

### tRNAseq: RNA isolation, tRNA library preparation and sequencing

Yeast total RNA was depleted from rRNA and sRNAs, followed by library preparation and Illumina sequencing. Yeast strains were grown in liquid cultures in SC-His medium until reaching OD_600_ of 0.9. Cells were harvested by centrifugation, washed once with water, with cell pellets corresponding to an OD_600_ of 9 OD subsequently frozen. Total RNA was then isolated from cell pellets using Epicentre Masterpure Yeast RNA Kit, which were processed with 4 times the volume of chemistry of the Epicentre kit; all remaining steps were done as described by the manufacturer (including DNaseI digest to remove residual DNA). 8 μg of isolated total RNA was separated from rRNA using the Illumina RiboZero Gold for Yeast rRNA removal kit, resulting in around 1.8 μg of rRNA-depleted RNA. sRNAs below 200 nt were subsequently removed using the Ambion MirVana miRNA Isolation kit (according to manufacturer’s instructions), resulting in around 300 ng of RNA. RNAs were checked for successful rRNA depletion and sRNA depletion using an Agilent Bioanalyzer RNA 6000 Nano. Illumina sequencing ready libraries were prepared using NEBNext Small RNA Library Prep Set for Illumina following the manufacturer’s instructions and checked on a Bioanalyzer DNA 1000. Illumina sequencing was performed on an Illumina NextSeq 500. tRNA quantification and differential expression testing Illumina sequencing data were quality controlled with FastQC v0.10.1 (de Sena Brandine and Smith, 2019). Sequencing adapters were trimmed with cutadapt v1.15 (Kechin *et al*., 2017). tRNAs were quantified at the transcript-level with Salmon v1.2.1 (Patro *et al*., 2017) using both pre-tRNA and mature tRNA transcript models. tRNAs were detected *de novo* using tRNAscan-SE v.2.0 (Lowe and Chan, 2016) on the tRNA neochromosome reference sequence. pre-tRNAs were modified with 10 nucleotide additional leader and trailer sequences based on the genomic coordinates. tRNAs derived from synthetic and native chromosomes were aggregated in a single reference transcriptome. The identical reference transcriptome was used to quantify each sample, independent of neochromosome presence, allowing for determination of misquantification rates on the neochromosome. Additional small RNA species were downloaded from Ensembl. Salmon was run with both sequence and position bias modelling enabled and in paired end mode. Differential expression testing was performed with Sleuth v0.30.0 (Pimentel *et al*., 2017).

### Replication timing profiles

Relative replication time was determined by sort-seq as described previously (Batrakou *et al*., 2020). Briefly, replicating (S phase) and non-replicating (G2 phase) cells were enriched from an asynchronously growing culture by FACS based on DNA content. As controls, G1 phase (α factor arrest) and G2 phase (nocodazole arrest) samples were also collected. For each sample, genomic DNA was extracted and subjected to Illumina sequencing to measure relative DNA copy number. Replication timing profiles were generated by normalizing the replicating (sorted S phase) sample read count to the non-replicating (sorted G2 phase) sample read count in 1 kb windows. The ratio was normalized to a baseline of one to control for differences in number of reads between the samples.

---

**Figure S1:**
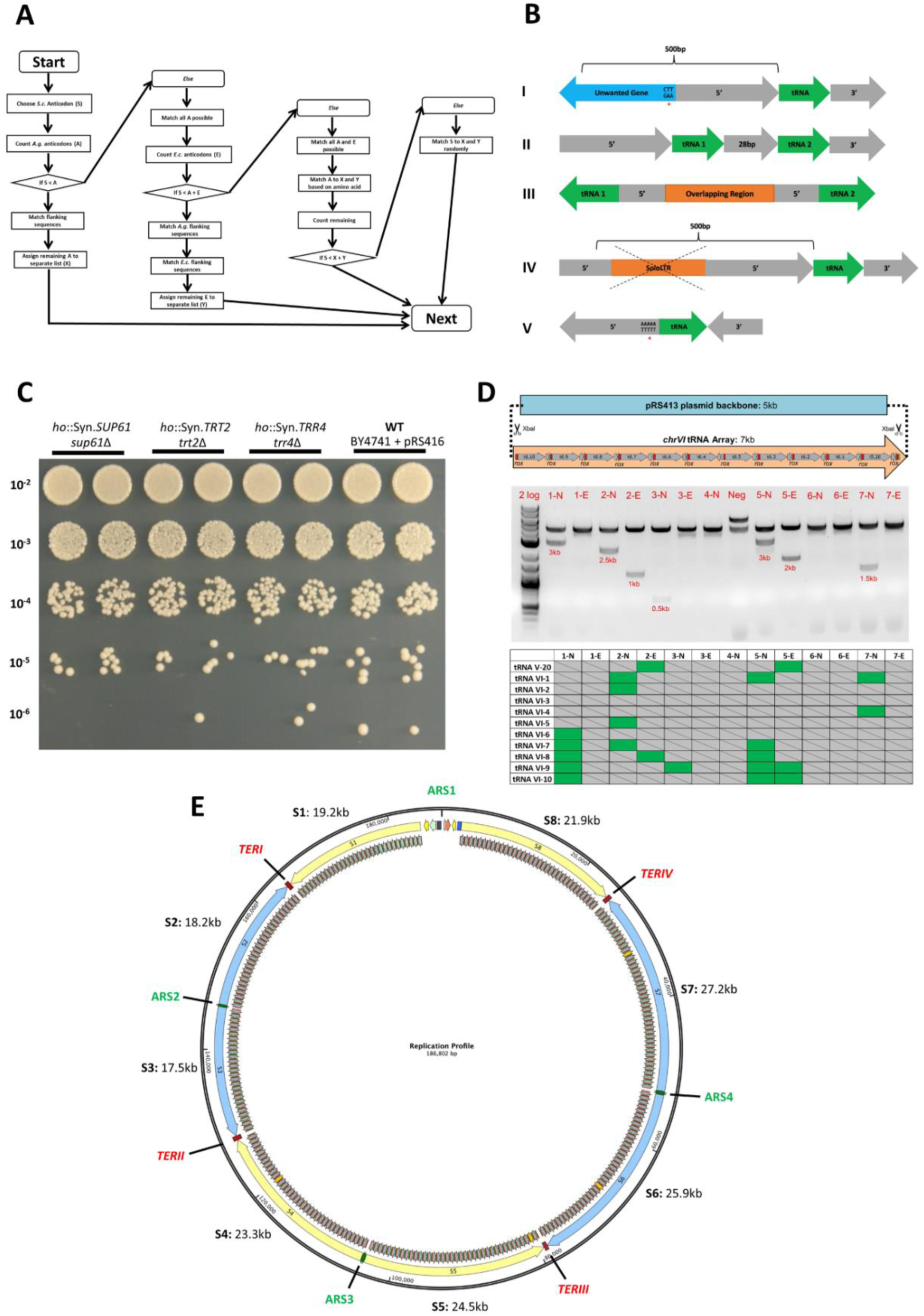
**(A) Flow chart of algorithm used to assign *S. cerevisiae* tDNAs to flanking sequences of *A. gossypii* and *E. cymbalariae***. The flow chart formed the basis of programming scripts used to generate the tRNA arrays of the tRNA neochromosome. The first stage of the algorithm identified a single anticodon family of *S. cerevisiae* (**S**) and assigned an *A. gossypii* (**A**) flanking sequences with the same anticodon based on the available pool. The remaining unassigned flanking sequences were then added to a separate list (**X**). For tDNAs that cannot be assigned *A. gossypii* flanking sequences based on this principle (**IF S>A**), the second stage of the algorithm performed the same process of assignment based on tRNA flanking sequences from *E. cymbalariae* (**E**) and unassigned flanking sequences were again added to a separate list (**Y**). For tRNA genes that cannot be assigned flanking sequences by matching anticodon from either of these two species (**IF S>A+E**), tDNAs were assigned flanking sequences based on matching amino acid from the two *A. gossypii* and *E. cymbalariae* lists (**X** and **Y**). Finally, the remaining tDNAs that cannot match any of these conditions were randomly assigned flanking sequences irrespective of anticodon, amino acid or species. **(B) Graphical representation depicting the removal/alteration of unwanted features in tDNA flanking sequences**. Otherwise, tDNAs and their associated flanking sequences were discarded from the available ‘pool’. (**I**) Unwanted genes in 5’ flanking sequence. Partially transcribed gene ‘starts’ identified in the 5’ flanking sequence were translationally silenced by altering the ‘ATG’ start codon to the ‘AAG’ lysine codon (denoted by the red asterisk). (**II**) Co-transcribed tDNAs separated by a 28 bp spacer region. The 5’ flanking sequence of tRNA 1 and the 3’ flanking sequence of tRNA 2 may be used (or otherwise discarded). (**III**) Shared 5’ flanking sequence between two divergent tDNAs. Only one of the two tRNA flanking sequences may be used based on relative anticodon requirement (or otherwise discarded). (**IV**) Solo LTR observed in 5’ flanking sequences of *E. cymbalariae*. The solo LTR was removed, with the 5’ flanking sequence extended to incorporate the full 500 bp. (**V**) Potentially misannotated tRNA orientation. This was inferred by the presence of a poly-thymidine residue in the 5’ flanking sequence (denoted by the red asterisk) and its absence in the 3’ flanking sequence. These tRNA flanking sequences were discarded from the available pool. **(C) Phenotypic colony morphology of each strain housing a synthetic copy of each essential tRNA gene.** The figure demonstrates that synthetic single-copy tRNA genes (*SUP61 (tS(CGA)C)*, *TRT2 (tT(CGU)K)* and *TRR4 (tR(CCG)L))* can complement the absence of each wild-type copy in BY4741 with no observable effect on colony size. Serial dilution and spotting of each strain (biological replicates) were undertaken onto SC-Ura. The dilution factor of each yeast sample is indicated on the left of the diagram. **(D) Dre recombinase-induced tRNA copy number adjustment: proof-of-concept. Upper diagram:** Illustrative map of the *chrVI* tRNA array on pRS413. The tRNA array consists of 10 tRNA genes from *chrVI* and one from *chrV*. *Rox* recombination sites are indicated in red. **Middle panel:** XbaI digestion pattern of plasmids recovered from BY4741 subjected to induction with Dre recombinase. The 5 kb band corresponds to pRS413, the second band of varying sizes corresponds to loss of tRNA genes on the array. Neg: negative control strain with tRNA array and empty pRS415 vector; N: non-induced; E: estradiol induced. **Bottom panel:** Sequencing analysis of isolates housing the *chrVI* tRNA array following post-induction with Dre recombinase. Green: tRNA gene remains; Grey boxes: tRNA gene is lost. The loss pattern of the tRNA genes indicates leaky expression of P_SCW11_-DreEBD. **(E) DNA replication profile of the tRNA neochromosome.** The tRNA neochromosome was divided into eight mega-arrays (S1 to S8; replication length in kb is denoted in the above figure), with each oriented away from the leading edge of the replication fork from four origins of replication (*ARS1*: *ARSH4*; *ARS2*: *chrF-444*; *ARS3*: *chrL-615*; *ARS4*: *chrM-794*). The latter three origins were obtained from *C. glabrata*. Yellow and blue arrows indicate orientation of tRNA genes and direction of replication fork. Replication termination sites (red; *TERI* to *TERIV*) were introduced at relatively even inter-origin distance to terminate replication at defined regions. The diagram was generated using SnapGene software (www.snapgene.com).

**Figure S2.**
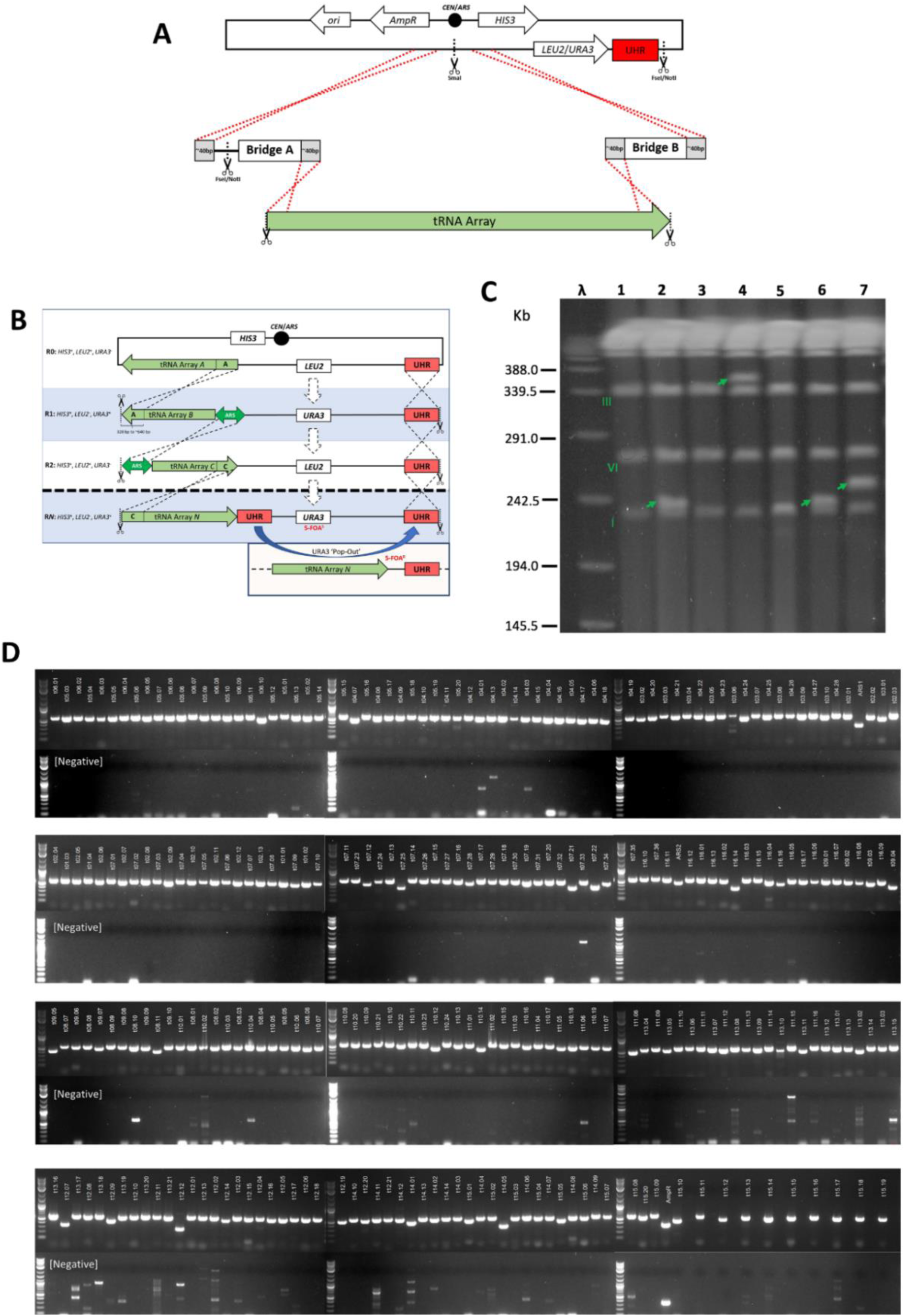
**(A) Introducing tRNA arrays into *LEU2* or *URA3* eSwAP-In vectors.** Two 330 bp to 650 bp ‘bridge’ sequences were amplified using PCR to facilitate construction using either Gibson Assembly or *in vivo* homologous recombination in yeast. Each bridge sequence was amplified using primers designed to contain the 40bp of homology required for assembly, together with a single FseI or NotI restriction cut site (Bridge A). Furthermore, these bridge sequences additionally served the purpose of providing homologous regions required for tRNA neochromosome construction (Bridge A was the last segment of DNA from the latter round of construction, Bridge B represents homology for a future round). Scissors with dotted lines represent restriction enzyme cut sites. The eSwAP-In vector contained a single SmaI restriction cut site to facilitate linearization. **(B)** Alternate representation of tRNA neochromosome construction and ‘pop-out’ of the *URA3* marker. The figure elaborates on the process of tRNA neochromosome construction. The black dot indicates the centromere, tRNA arrays are indicated by colored arrows, auxotrophic markers by green arrows and the 500 bp UHR with a red box. The FseI/NotI restriction sites are indicated by scissors and black dotted lines. During Round 1, the ‘A’ region of tRNA Array B facilitates homologous recombination, with the *LEU2* marker replaced with the *URA3* marker. For all future rounds, marker swapping may be facilitated with any region of homology (*e.g.* ARS elements as denoted in the above figure), with the UHR remaining static. To induce *URA3* pop-out (bottom panel), a second UHR is introduced to facilitate homologous recombination. *URA3* converts 5-FOA into the toxic 5-fluorouracil – only cells that remove functionality of the URA3 gene will survive. **(C) Pulsed-field gel pattern of strains housing linear and circular variants of the tRNA neochromosome constructed in BY4741**. The above gel image shows an unexpected heterogeneity in tRNA neochromosome size. Lanes 1 and 3 are strains housing circular variants, lanes 2 and 6 are replicate isolates (n = 2) of strains housing a neochromosome linearized at *TERIII* (∼240 kb band), lanes 4 and 7 are replicate isolates of strains housing a neochromosome linearized at *TERII* (∼350 kb and ∼250 kb bands, respectively) and lane 5 is a wild-type empty plasmid control. The expected size of the tRNA neochromosome is ∼187 kb. Wild-type yeast chromosomes are indicated by Roman numerals and bands corresponding to the neochromosome are indicated by green arrows. **(D) Representative PCR tag analysis of the tRNA neochromosome.** PCR tag names are denoted at the top (a full list may be found in Table S4). The negative control (bottom panels) was undertaken on BY4741 + pRS413. ‘AmpR’ denotes a positive control designed to amplify *bla* on pRS413.

**Figure S3.**
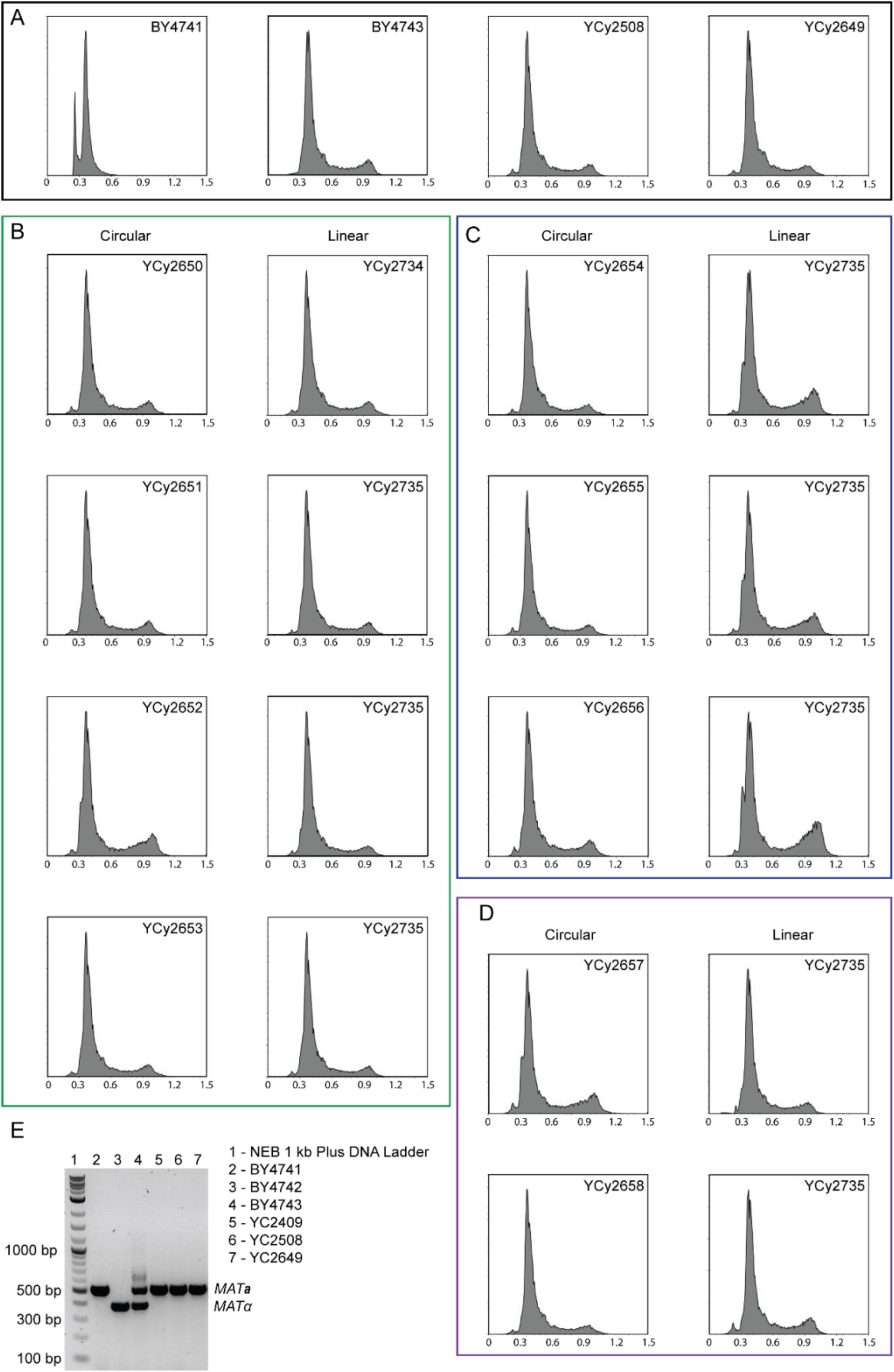
Ploidy and mating type determination of tRNA neochromosome strains. **(A)** DNA content analysis of haploid BY4741 and the diploid strains BY4743, YCy2508 and YCy2649 serving as controls for panels B, C and D. **(B-D)** DNA content analysis to determine the ploidy of strains pre- and post-linearization, all strains show the pattern for diploid DNA content. **(B)** Strains linearized at the *TER* sites. **(C)** Strains linearized at *ARS* sequences. The strains show a higher proportion of cells with 4n DNA content in contrast to all other strains. However, this seems not to influence the phenotype of the strains under the tested conditions (*cf*. Fig. S6). **(D)** Strains linearized close to *CEN*. (A-D) 100,000 total events were detected and each histogram consists of minimum 70,000 gated events. Y-axis displays number of events and x-axis intensity (AU x 1,000). **(E)** Multiplexed *MAT*-locus PCR according to Harari *et al*. (2018b) with universal-(5’-AGTCACATCAAGATCGTTTATG-3’), *MAT**a***-specific (5’-ACTCCACTTCAAGTAAGAGTTTG-3’; ∼500 bps) and *MATα*-specific primers (5’-GCACGGAATATGGGACTACTT-3’; ∼400 bps). Analysis confirms that the final tRNA neochromosome strain is *MAT**a***. Taking the results together, as well as the ability of the strains to mate with *MATα*-strains (data not shown), suggest a whole-genome duplication resulting in a *MAT**a**/**a*** genotype.

**Figure S4.**
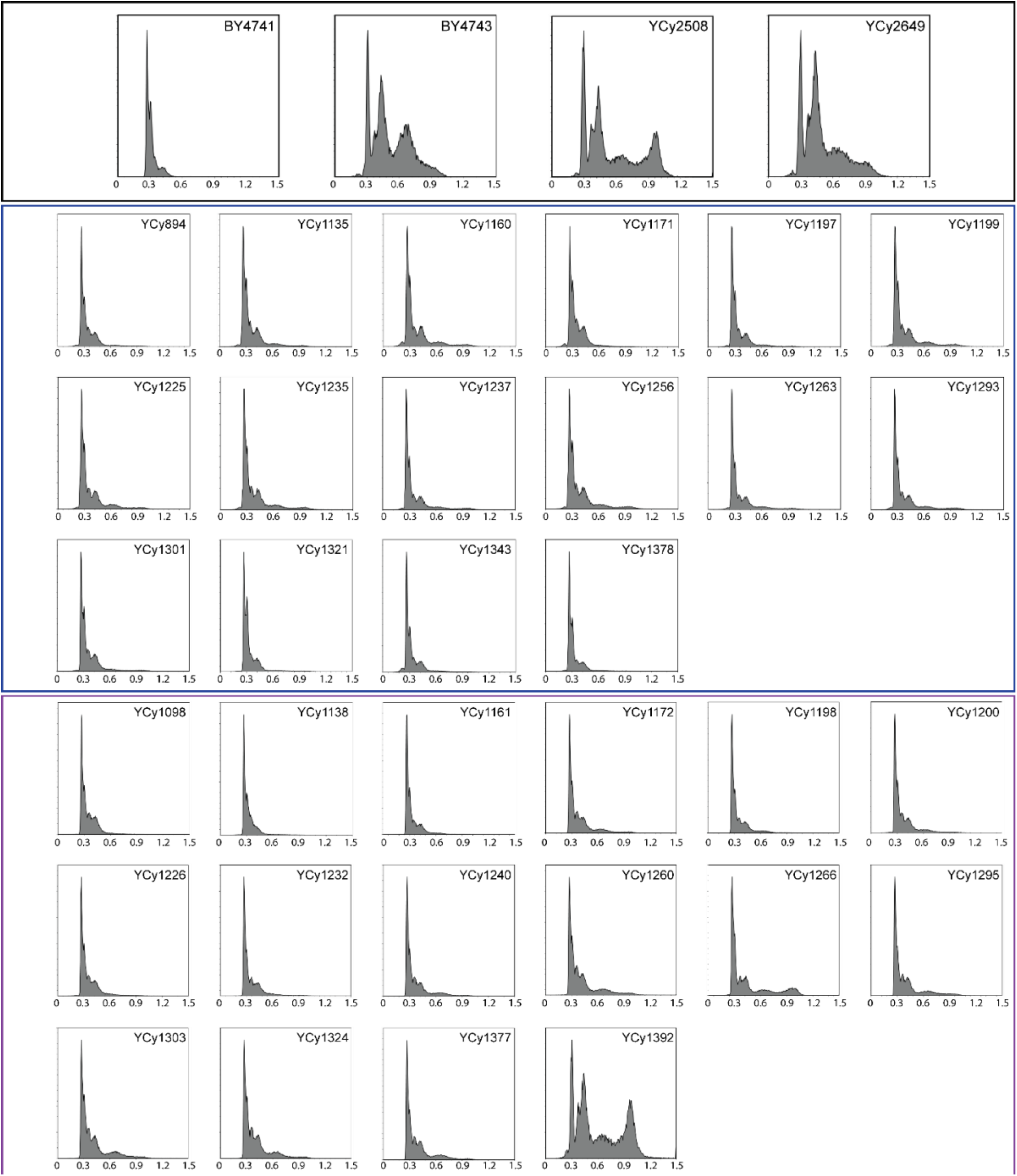
Ploidy determination of individual intermediary strains following each round of eSwAP-In. DNA content was determined by flow cytometry for individual strains in comparison to BY4741, BY4743, YCy2508 and YCy2649. All surveyed strains except YCy1392 are haploid. YCy1392 presumably underwent a whole genome duplication independent of spheroplast transformation. The upper colored frames correspond to controls (black), the middle frame represents consecutive eSwAP-In rounds of all tRNA arrays in BY4741 (blue) and the bottom panel corresponds to YLM896 (or Syn III/VI/IXR; purple). 100,000 total events were detected for each sample and each histogram consists of minimum 50,000 events. Y-axis displays number of events and x-axis intensity (AU x 1,000).

**Figure S5.**
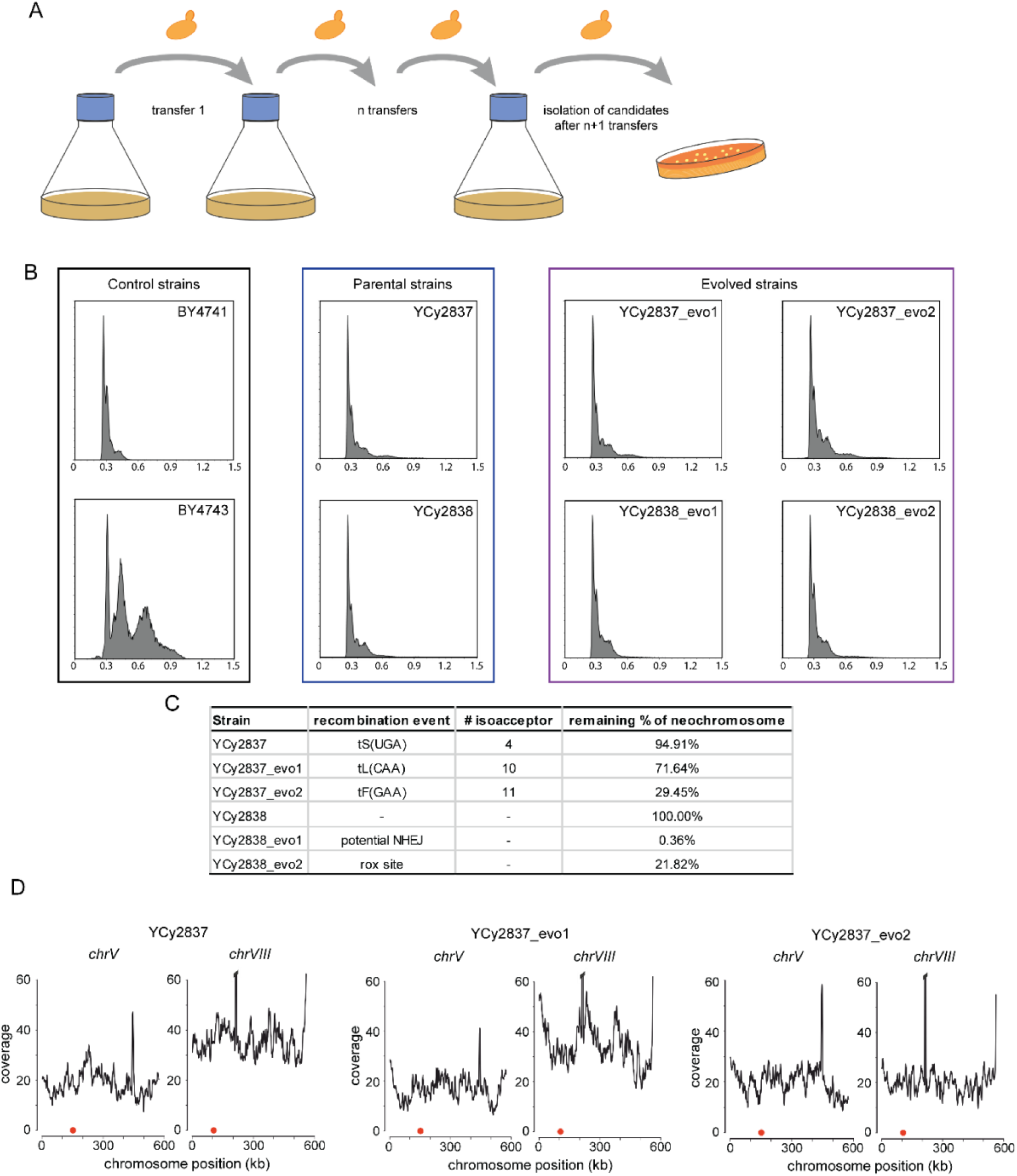
Time course experiment for tRNA neochromosome stability, set-up and results. **(A)** The time course experiment was performed as batch transfer where yeast cultures were back-diluted every 24 h to an OD_600_ of 0.1. Single isolates were obtained after serial dilution and plated onto SC-His medium. **(B)** Flow cytometry analysis of DNA content of the starting strains and four evolved isolates in comparison to BY4741 and BY4743. All tested strains are haploid. 100,000 total events were detected for each sample and each histogram consists of minimum 50,000 events. Y-axis displays number of events and x-axis intensity. **(C)** Structural variations in the tRNA neochromosome detected by Nanopore Sequencing of the parental strains and evolved isolates. **(D)** Nanopore Sequencing revealed an aneuploidy for *chrVIII* in the parental strain and the evolved strain containing >70% of the tRNA neochromosome content while the aneuploidy is lost in the strain only containing 30% of the tRNA neochromosome content. Visualization of c*hrV* serves as a reference for the average genome coverage in each sequencing experiment. Red dot indicates position of *CEN*, *CUP1-1*/*CUP1-2* amplification in *chrVIII* exceeds y-axis scale and data is omitted for visualization purpose.

**Figure S6.**
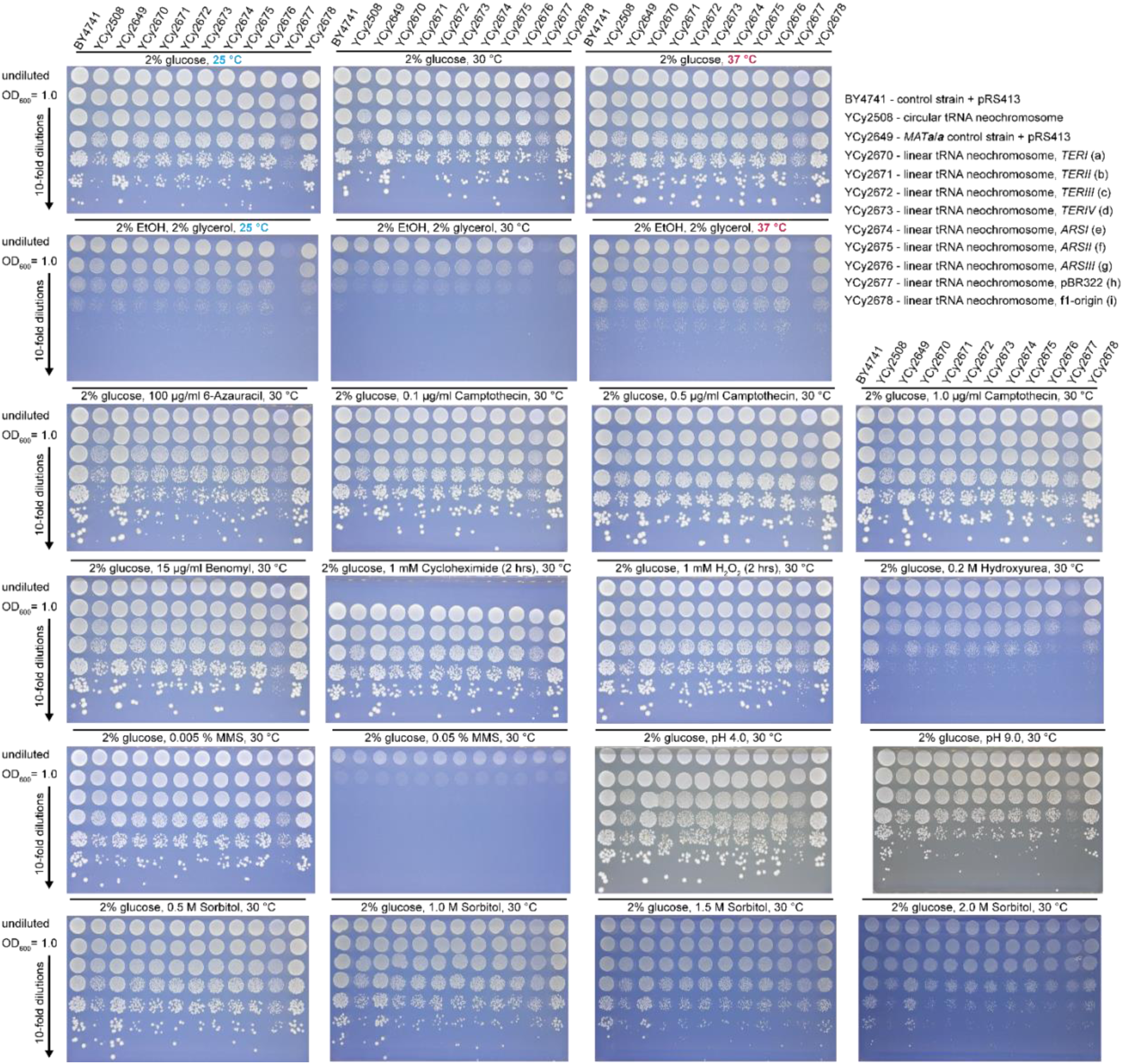
Phenotyping tRNA neochromosome derivatives across a range of 22 growth conditions. Each panel shows a single media condition with identical strain distribution of one circular and nine linear derivatives. In each panel, the top row represents the undiluted overnight culture followed by a 10-fold serial dilution from a culture normalized to an OD_600_ of 1.0 (second row and below). Similar growth patterns can be observed throughout each assay, except for 6-azauracil and 0.2 M hydroxurea where the growth deficiency of the tRNA neochromosome strains increases. The strain YCy2677 was later revealed to be a mitochondrially deficient rho^-^ mutant (indicated by a lack of growth on non-fermentable carbon sources). All spotting assays where performed on SC-His at 30 °C for 72 hrs unless specified otherwise. Carbon source and alteration are stated above each panel. *cf*. Figure 3A for visualization of linearization sites.

**Figure S7.**
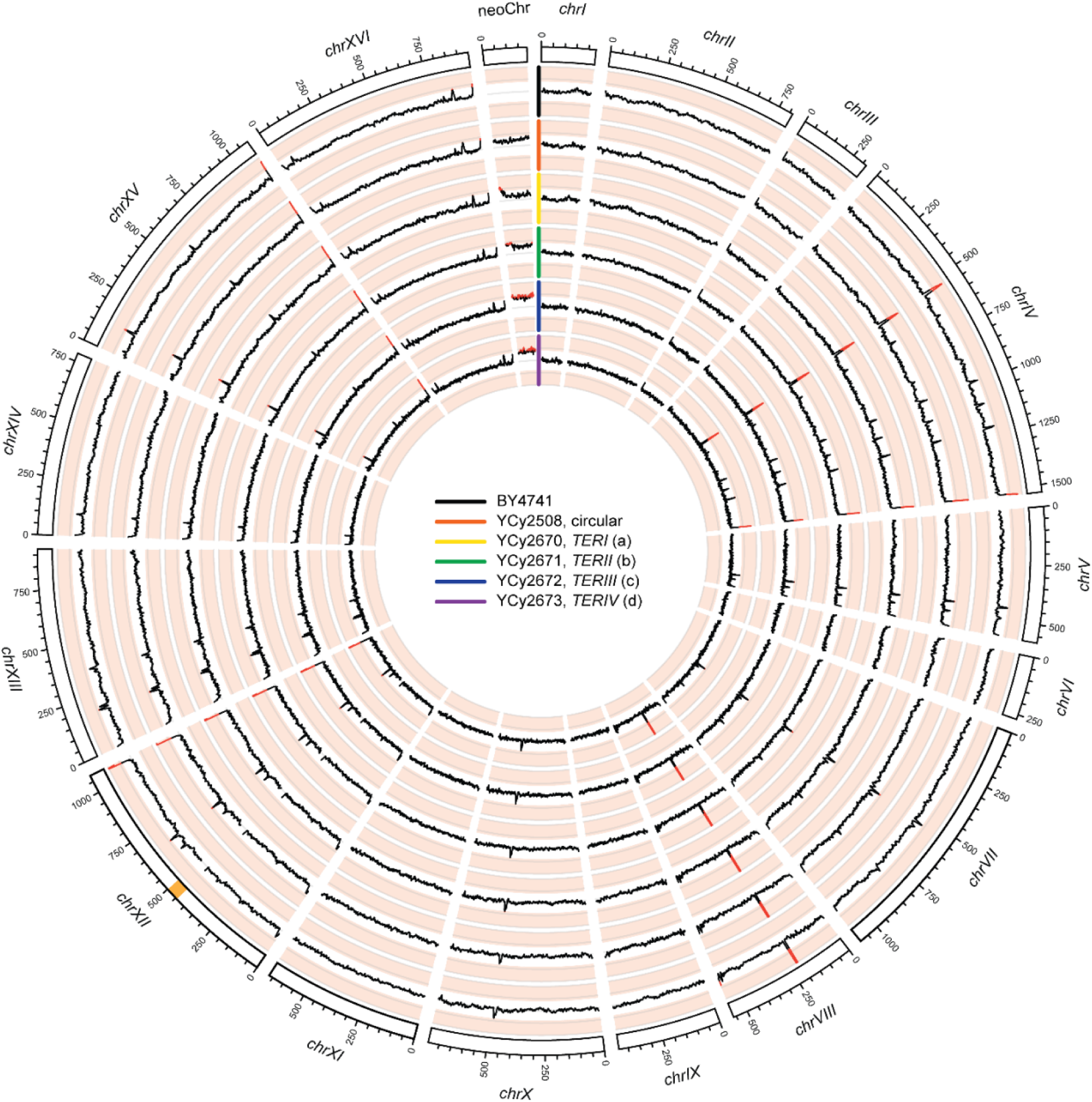
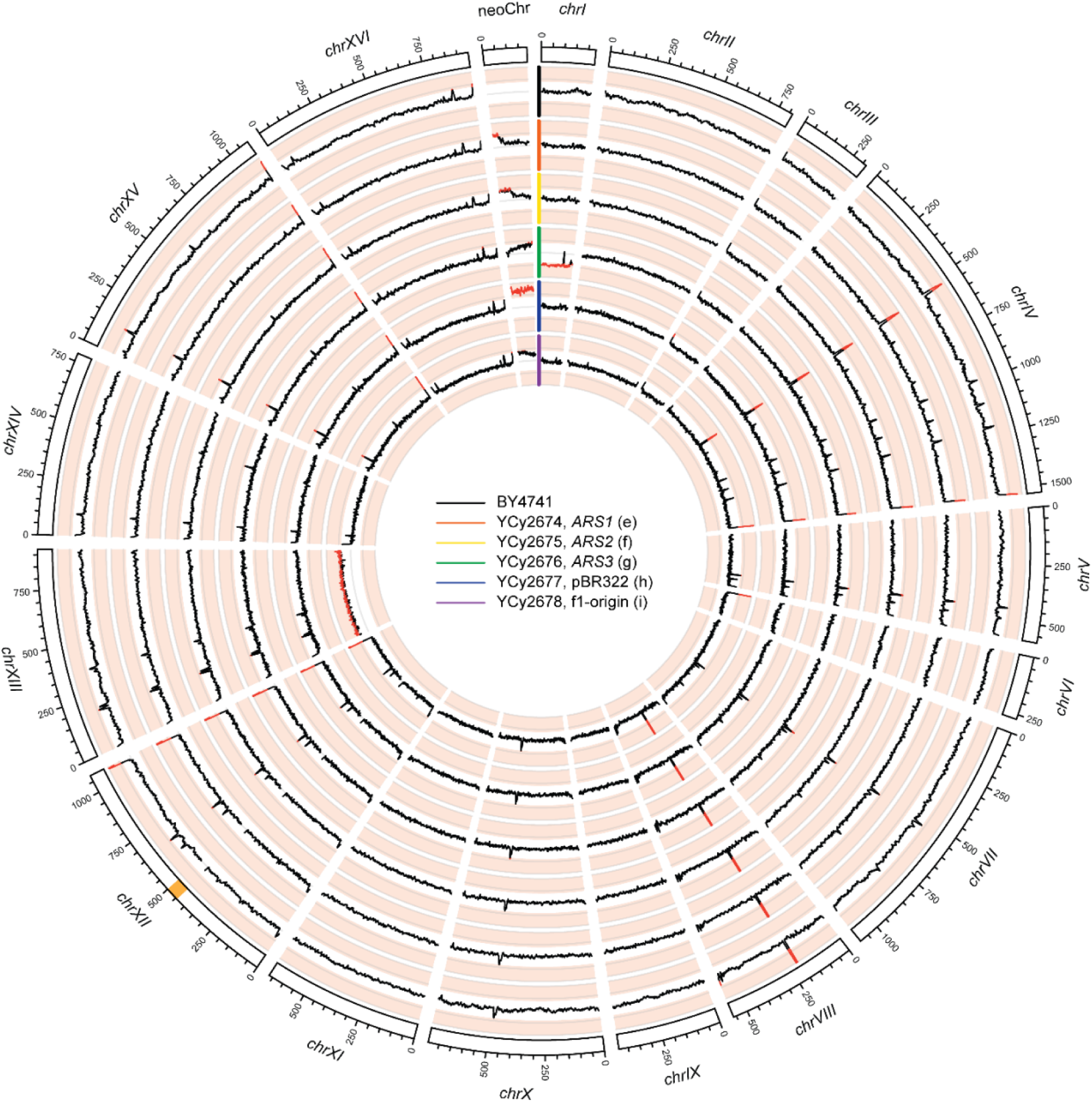
Short-read sequencing data for all linearized tRNA neochromosome variants. The above Circos plots indicate sequence read coverage for tRNA neochromosome variants obtained from stationary phase cells in comparison to BY4741. Linearization locations are indicated in the center of each circle, with different colors denoting each strain. The sequencing coverage was normalized within each sample to visualize aneuploidies, corresponding to ≥ 1.5 and < 0.5-fold coverage (scale from 0 to 2; indicated by red coverage lines for threshold passing). A moving window average function (5,000 bp average 500 bp steps) was applied to smoothen the curves using a custom R script. The rDNA region, indicated orange in chromosome XII, was excluded from plotting. In both plots the outer most ring is the control strain BY4741. Coverage analysis indicate a −1 aneupoidy for *chrI* in YCy2676 and a +1 aneuploidy for *chrXIII* in YCy2677. Notably, the strains containing the tRNA neochromosome linearized at suboptimal positions (YCy2674 to YCy2678) show alterations in tRNA neochromosome coverage, however no size alteration were detected during size analysis (*cf*. Figure 3). There are no segmental coverage alterations specific for an individual strain. Observed peaks correspond to known variations such as *CUP1-1*/*CUP1-2* in *chrVIII*, known to have variable number of tandem repeats and can further be present as extrachromosomal circular DNA (eccDNA). Plots are generated using Circos (Krzywinski *et al*., 2009). Sequencing data is available at Bioproject PRJNA884518.

**Figure S8.**
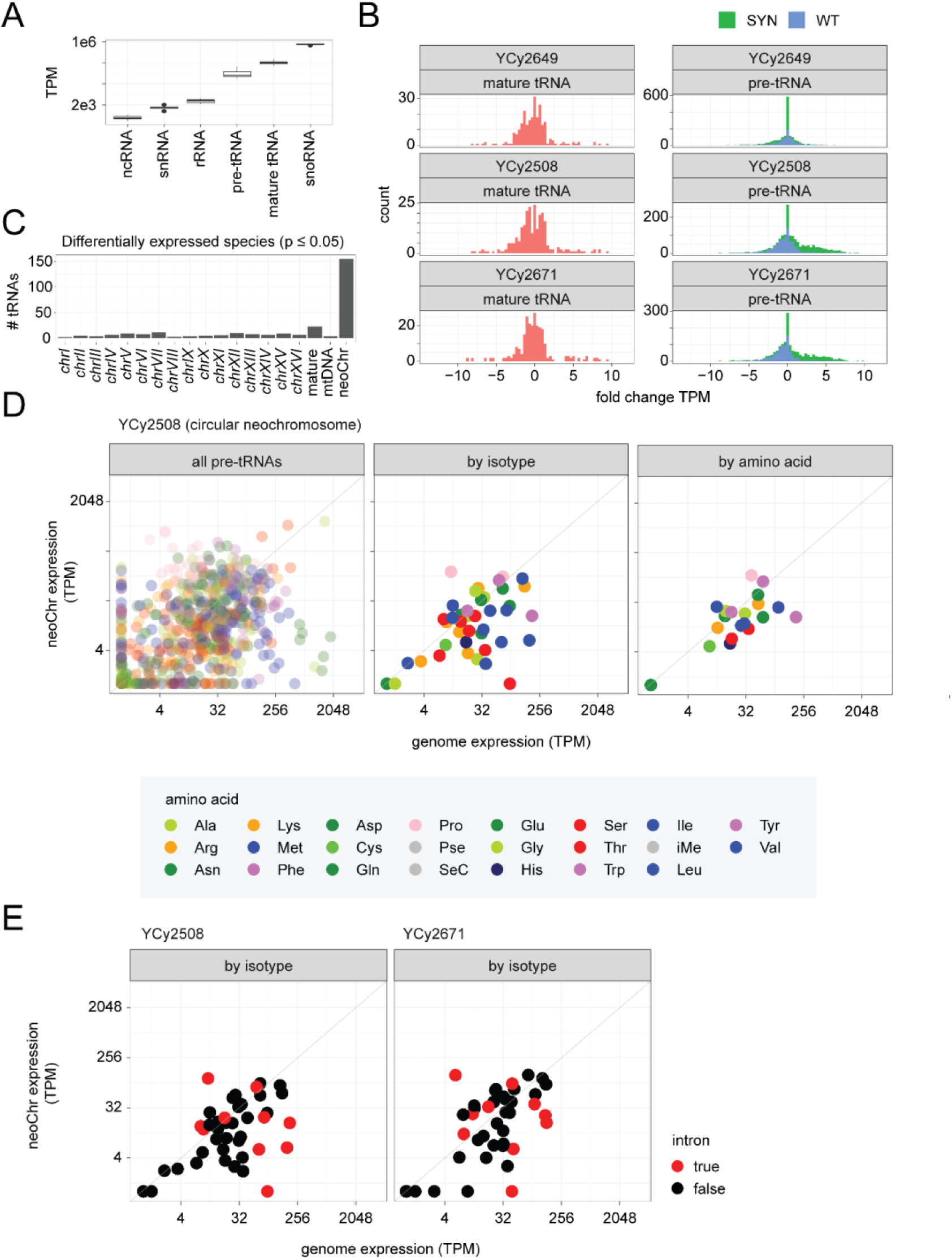
tRNA sequencing of circular and linear tRNA neochromosome variants. The figure displays results for YCy2508 (circular tRNA neochromosome), YCy2649 (BY4741 + pRS413 control) and YCy2671 (linearized tRNA neochromosome). **(A)** Quantification of all sequenced small RNA species. **(B)** Distribution of expression fold changes measured for mature and pre-RNA species originating either from the synthetic tRNA neochromosome or the native genome in a strain without the neochromosome (YCy2649) and strains with either a circular version of the neochromosome (YCy2508) or a linear version (YCy2671). **(C)** Number of differentially expressed tRNA species on each chromosome (p <= 0.05). **(D)** Pairwise pre-tRNA expression levels for all pre-tRNAs originating from the neochromosome compared to levels measured from the native loci they were designed to replace (left). Alternatively, pre-tRNA expression levels were aggregated and averaged (median) by isotype (center) or amino acid (amino acid). Pearson correlation coefficient (r) reported for each association. Colors denote the amino acid with which the tRNA species is charged. **(E)** As in D, with colors corresponding to the presence or absence of introns. TPM: transcripts per million.

**Figure S9.**
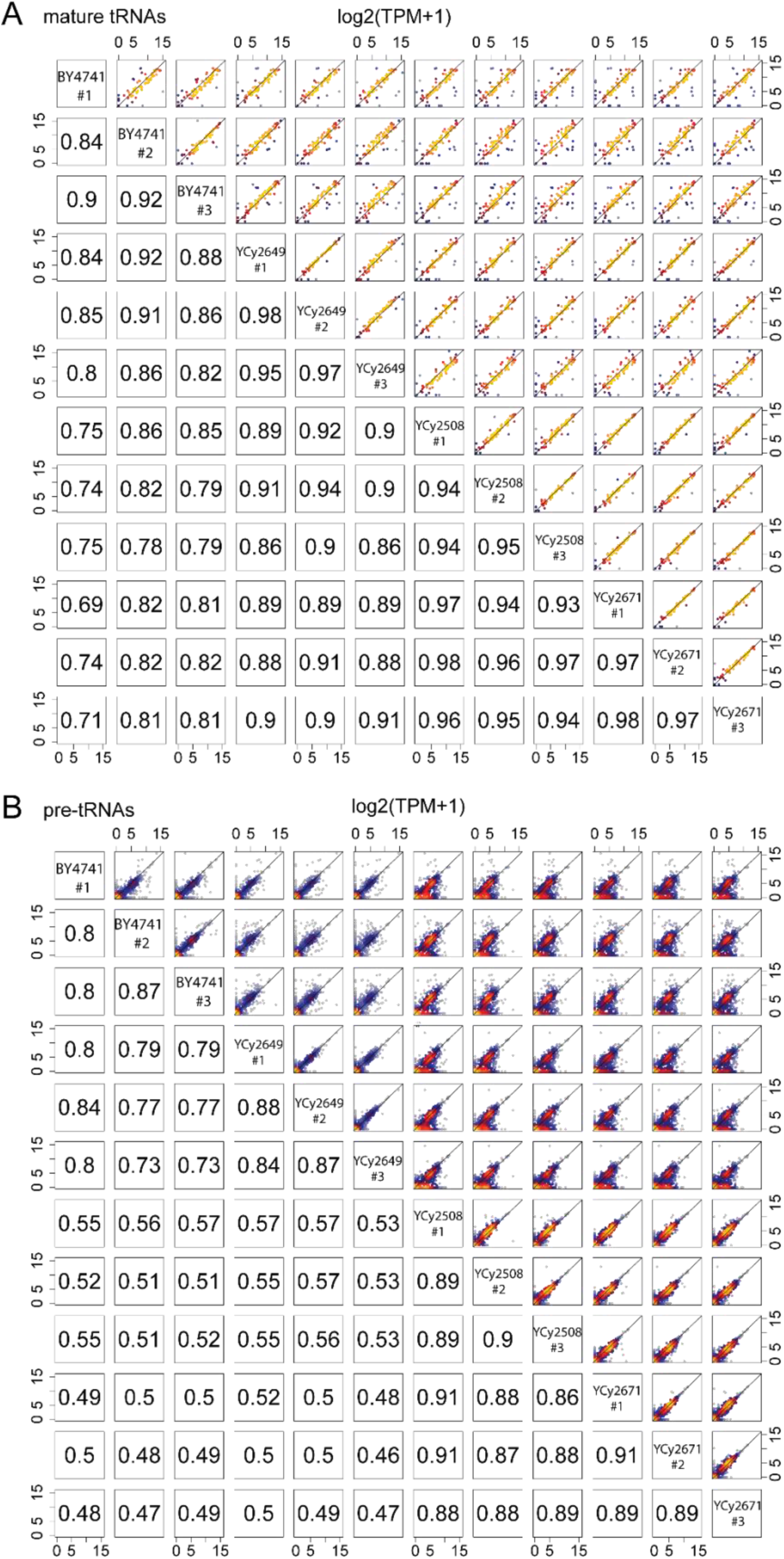
Comparative tRNA expression between sample replicates. Strain IDs are indicated in the middle of each chart. **(A)** Pairwise mature tRNA expression levels across all strains and replicates. **(B)** Pairwise pre-tRNA expression levels across all strain and replicates. Numeric values denote Pearson correlation coefficient (r). TPM: transcripts per million.

**Figure S10.**
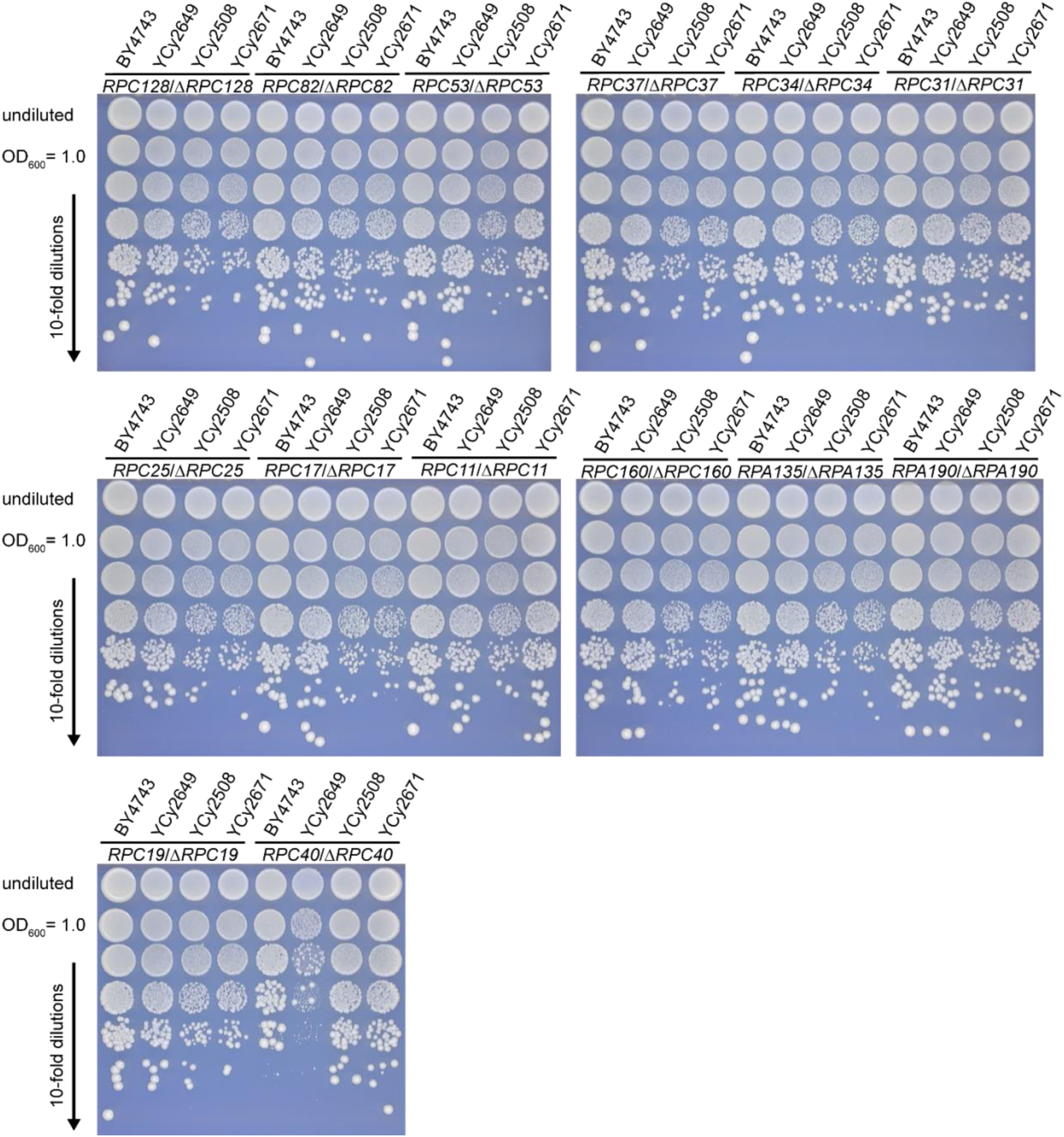
RNAPIII and RNAPI specific and shared subunit deletion. In the above figure, heterozygous diploids were generated with single subunit deletions of RNAPIII, RNAPI or shared subunits. Growth media conditions are SC-His, 2% glucose at 30°C for 72 hrs. In each panel, the top row represents the undiluted overnight culture followed by a 10-fold serial dilution from a culture normalized to an OD_600_ of 1.0 (second row and below). BY4743 is the wild-type diploid control, YCy2649 is the *MAT***a**/**a** control, YCy2508 is the homozygous diploid housing a circular tRNA neochromosome and YCy2671 is a homozygous diploid strain containing a linear variant of the tRNA neochromosome.

**Figure S11.**
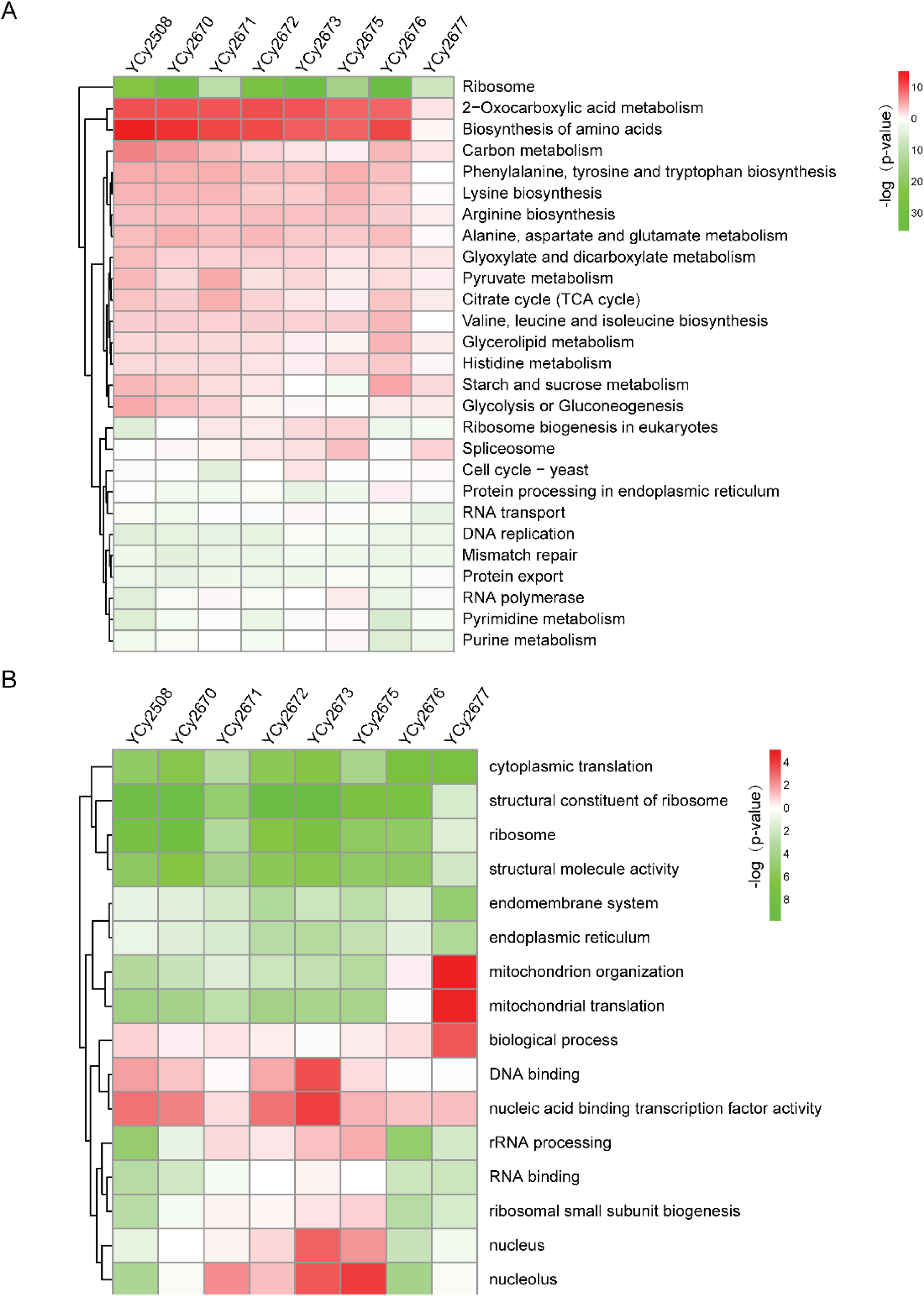
RNAseq data visualization for all analyzed tRNA neochromosome strains. Differential expression of circular tRNA neochromosome strains normalized to the homozygous diploid control strain (YCy2649). **(A)** KEGG and **(B)** GO analysis demonstrate a general upregulation of amino acid biosynthesis and a down regulation of translational processes. YCy2677 is the rho-strain which explains the upregulation of mitochondrial processes. Up-regulated and down-regulated features are labeled in red and green, respectively.

**Figure S12.**
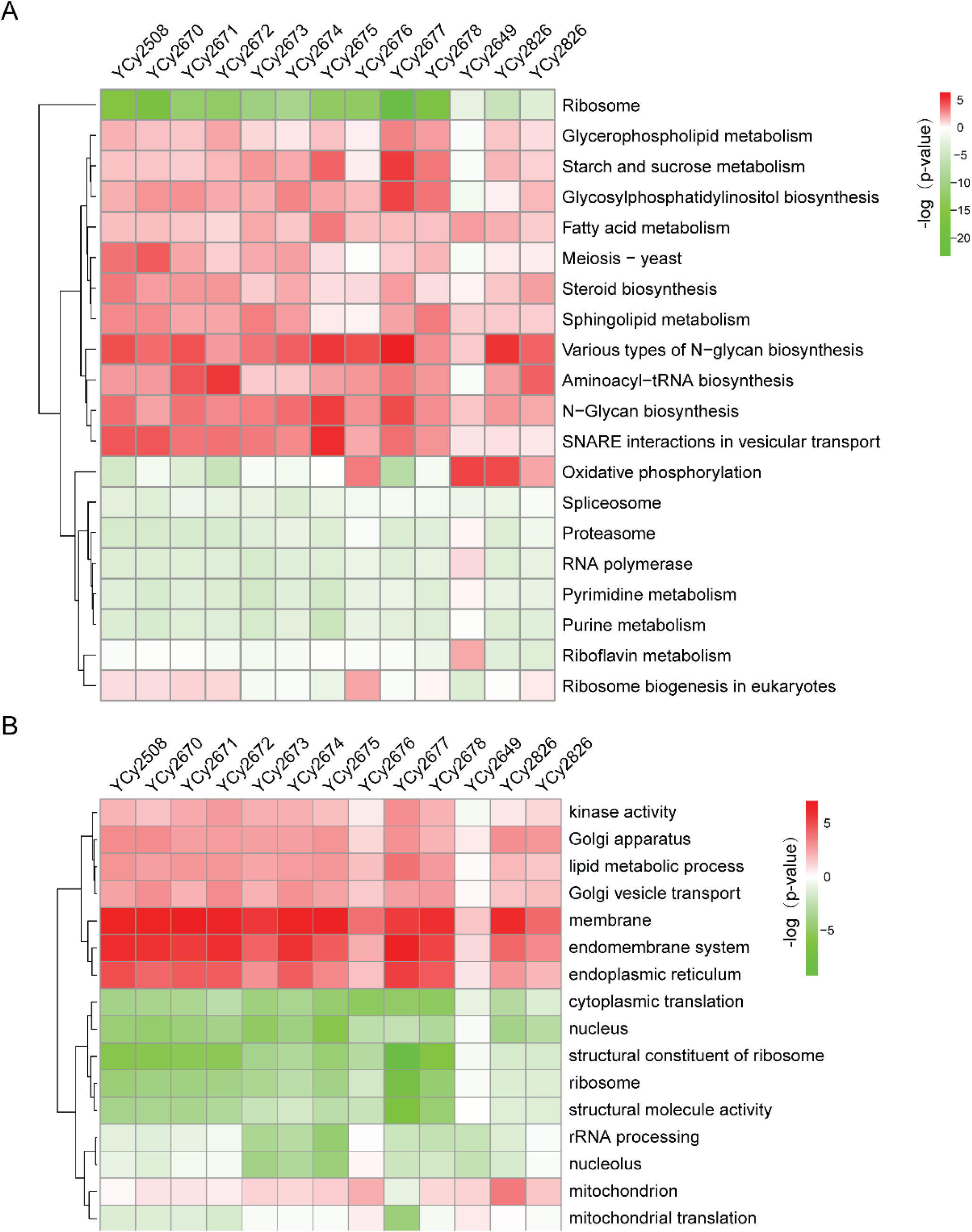
Proteome data visualization for all analyzed tRNA neochromosome strains. Proteomic analysis of the tRNA neochromosome strains normalized to the homozygous diploid control strain (YCy2649). **(A)** KEGG and **(B)** GO analysis globally show a similar pattern to that of transcriptomics. However, an additional increase in membrane-associated proteins and endoplasmic reticulum can be observed. Up-regulated and down-regulated features are labeled in red and green, respectively.

**Figure S13.**
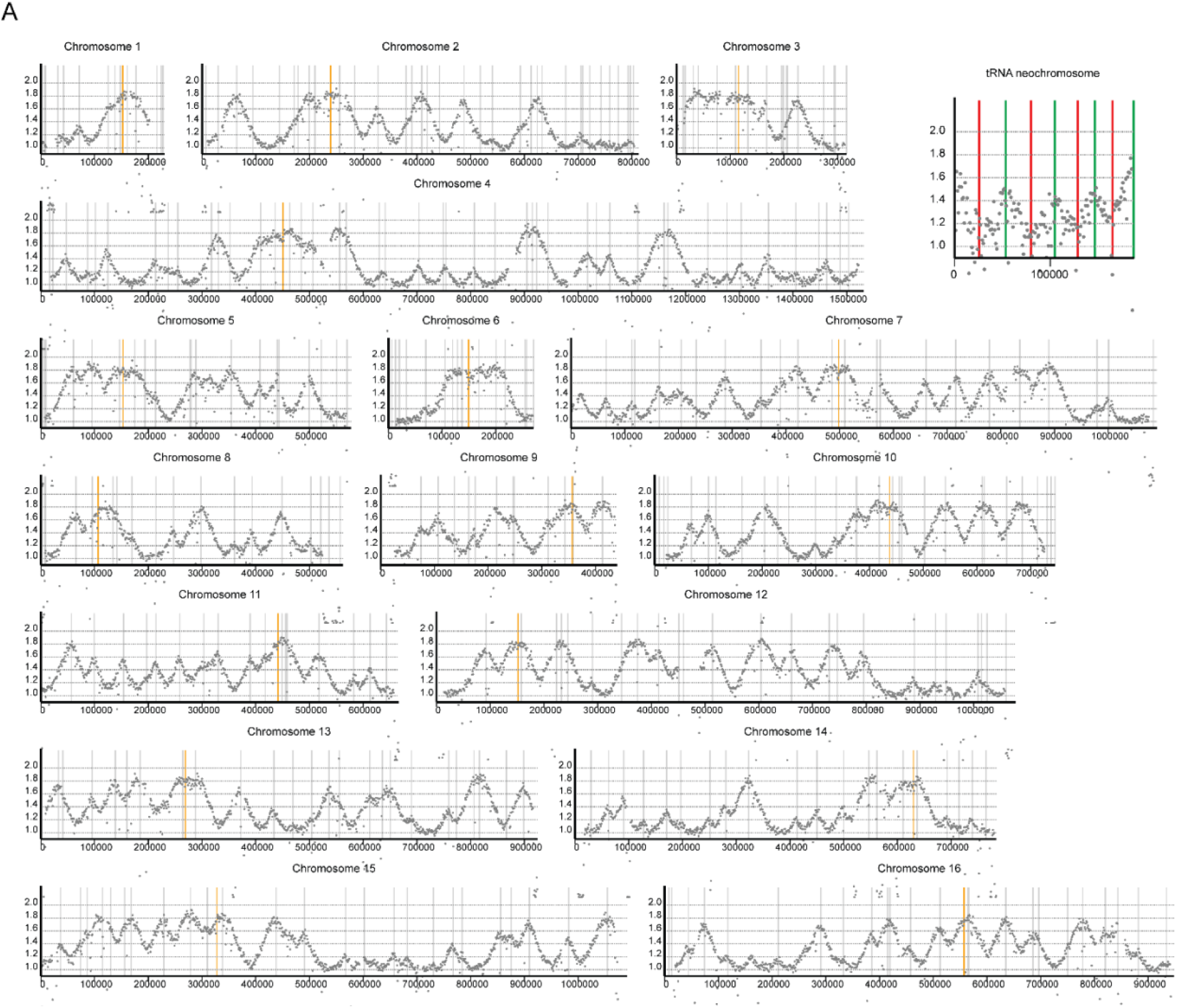
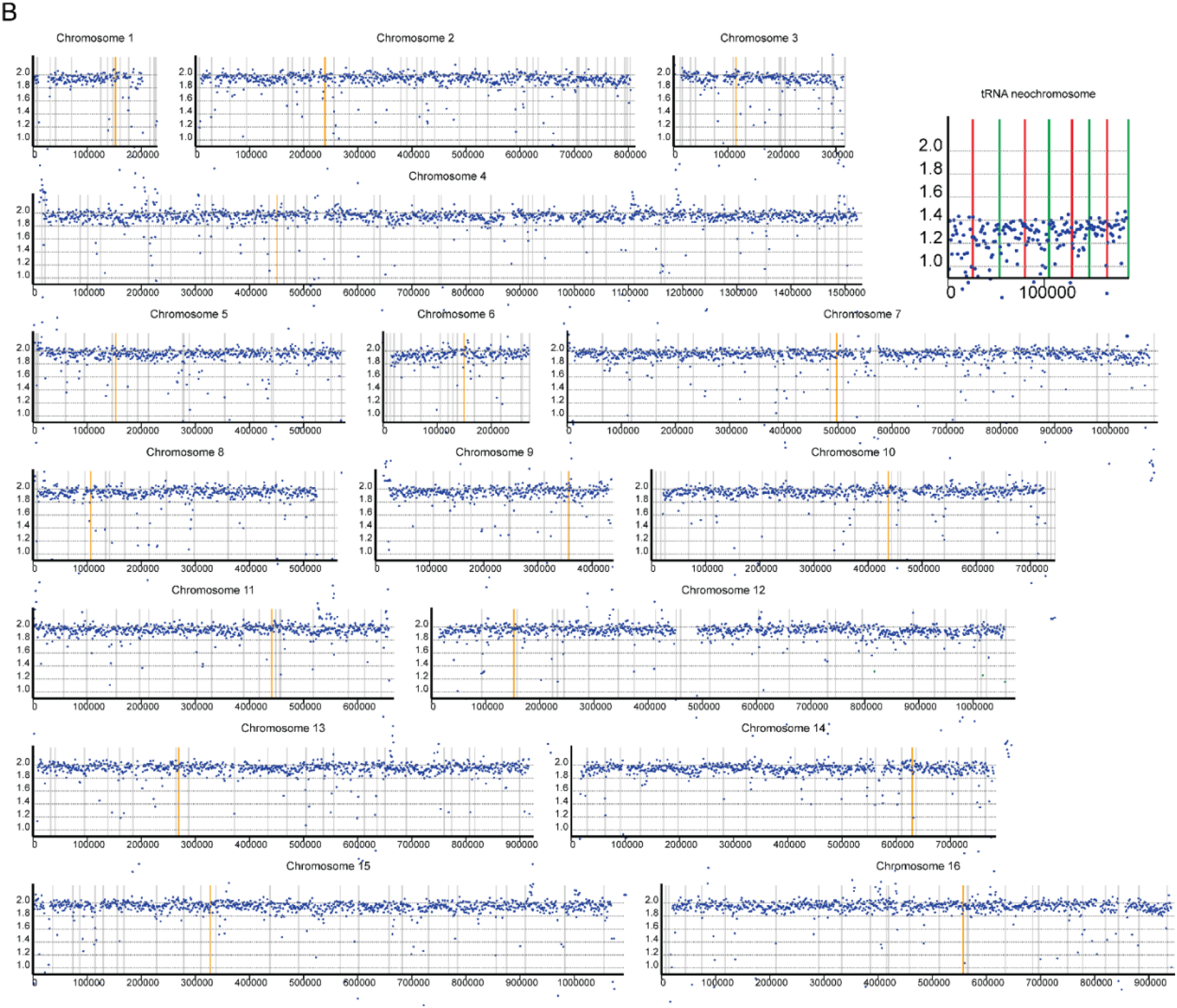
DNA replication profiling of S phase and G1 phase cells in a circular tRNA neochromosome variant (YCy2508). (A to C) *ARS* (gray or green), *CEN* (orange), *TERI-IV* (red). **(A)** Replication pattern of S-phase cells. Peaks correspond to origins of replication and pattern is indistinguishable from strains without the tRNA neochromosome (Müller and Nieduszynski, 2012). **(B)** Replication pattern of α-factor arrested cells (G1). All chromosomes show a normalized read-abundance of ∼2-fold while the tRNA neochromosome shows a lower abundance of ∼1.4-fold.

